# A genetic algorithm for self-supervised models of oscillatory neurodynamics

**DOI:** 10.1101/2024.12.31.630823

**Authors:** Hamed Nejat, Jason Sherfey, André M. Bastos

## Abstract

Predictive processing theories propose that the brain builds internal models of its environment by reducing the discrepancy between internally generated predictions and external sensory signals. Prior work has linked these processes to oscillatory activity in gamma (40-100 Hz) and alpha/beta (10-30 Hz) frequency ranges. Current computational approaches face a trade-off: abstract predictive-processing models can implement self-supervised computations but often omit oscillatory spiking dynamics, whereas biophysically constrained spiking models can generate neural rhythms but often require extensive manual tuning. Here, we introduce the Genetic Stochastic Delta Rule (GSDR), an evolutionary optimization framework for fitting nonlinear neural models to electrophysiological objectives. We first evaluate GSDR in simplified optimization settings, then apply it to spiking-network objectives involving firing rates, beta/gamma spectral ratios, and empirical macaque stimulus-evoked gamma dynamics from visual cortex. We show that GSDR can search constrained synaptic parameter spaces, reduce reliance on manual tuning, and reproduce spectral and circuit-level phenotypes associated with predictive routing. We also used Izhikevich simulations as a model-class robustness analysis, showing that the approach is not limited to the original Hodgkin-Huxley-style implementation. These results position GSDR as a methodological framework for multi-objective exploration of oscillatory neural models.

**Author summary:** In predictive processing theories, the brain is hypothesized to build internal models of its environment. Empirical and theoretical studies suggest that neuronal oscillations are important components of this process, and abnormal oscillations are also linked to disorders such as schizophrenia. To study such mechanisms, computational neuroscience needs models that can express biologically meaningful spiking and oscillatory dynamics while also being trainable without extensive manual tuning. We developed the Genetic Stochastic Delta Rule (GSDR), a self-supervised evolutionary optimization framework for fitting nonlinear neural models to objectives. GSDR combines objective-guided search, stochastic exploration, genetic selection/deselection, and an activity-dependent MCDP update term. We show that GSDR can tune spiking networks toward beta/gamma spectral objectives and empirical stimulus-evoked gamma dynamics. The results do not prove predictive routing or identify a unique biological circuit; rather, they show that GSDR can identify candidate circuit configurations and can generalize beyond the original Hodgkin-Huxley-style model to Izhikevich simulations.

## Introduction

Understanding the brain requires linking theoretical mechanisms of neurons to empirical observations in vivo. Biophysical neural models in neuroscience [1–6] help bridge fundamental neuronal properties, neurophysiology, cognition, and behavior. Models differ in their level of abstraction: conductance-based models can expose channel- and receptor-level mechanisms, population models can preserve efficient cell-level dynamics, and population or neural-mass models can summarize large-scale dynamics. Hodgkin-Huxley-style models [11,12] are useful when the scientific question concerns ionic currents, conductances, or their voltage and time dependencies. These modeling choices make it possible to ask how changes in ion-channel properties [13–15], cell types [16–18], cortical layers [19–21], neurotransmitters [22–24], neuronal metabolism [25–28], oscillations [29–32], and synaptic integration [32–34] affect perception, cognition, and behavior.

Models of neuronal oscillations are particularly important for predictive processing theories [35–41], which propose that top-down and bottom-up processing are associated with distinct oscillatory channels. Top-down processing reflects internal or cognitive state variables such as attention or prediction [42,43], whereas bottom-up processing reflects how external sensory states are signaled [44–46]. Predictive routing (PR) [47] is one theory that links predictive processing to empirically observed oscillatory dynamics. PR proposes that gamma-band activity and increased spiking carries bottom-up signals from lower-order sensory cortex toward higher-order areas, whereas lower-frequency alpha/beta activity reflects top-down predictions that modulate sensory processing [47,50]. In the present manuscript, PR is used as a motivating neuroscience case study because its proposed building blocks include spiking dynamics and beta/gamma state changes. The goal is not to prove PR or to implement a full predictive task; instead, we test whether automated optimization can fit spiking neural models to spectral and empirical objectives relevant to this framework.

A mechanistic account of how these oscillatory dynamics emerge from neuronal circuits, and how they might support predictive routing, remains incomplete. Numerous studies have modeled oscillatory interactions with conductance-based, and population-level approaches [17,18,42,53–57]. These models can generate oscillatory dynamics, including stimulus-evoked gamma oscillations. Many use Pyramidal-Interneuron networks (PIN), consisting of input drive to pyramidal neurons and feedback inhibition from interneurons. The oscillation frequency depends on inhibitory synaptic time constants, recurrent connectivity, and input structure, such that a network can be manually tuned to express PIN-gamma (PING) or PIN-beta (PINB). Networks of interconnected inhibitory neurons can also generate gamma oscillations without pyramidal participation, an alternative mechanism referred to as Interneuron Network Gamma (ING) [33]. In both PING and ING, synaptic connectivity, internal noise, and bottom-up inputs can generate rhythms that are either highly synchronous (strong PING, [1,2,17,53]) or sparse and irregular across neurons (weak PING, [58–61]). Biologically observed dynamics in healthy brains are often more consistent with the latter sparse and irregular regime.

A practical limitation of many nonlinear spiking models is that they require extensive manual tuning. Exhaustive parameter sweeps can become computationally intractable, especially when models include many synaptic weights, time constants, input gains, or cell classes. Some optimization methods have been applied to neural models [62–64], but many approaches still require strongly hand-specified parameter choices or do not include activity-dependent terms. We therefore frame the present work around self-supervised optimization in a specific sense: models do not rely only on external labels or experimenter-provided target updates. Instead, learning can combine objectives with structure in the model activity itself. In this manuscript, the activity-dependent component is mutual-correlation dependent plasticity (MCDP), which uses membrane-potential similarity over time to drive updates within the existing model connectivity.

In this work, we introduce the Genetic Stochastic Delta Rule (GSDR), a parameter optimization framework for biophysically constrained spiking models using evolutionary algorithms [69–72]. GSDR is inspired by the stochastic delta rule [73–75] and genetic algorithms [76–78]. In its basic form, GSDR is a multi-variable optimization method for exploring nonlinear parameter spaces to optimize one or more objectives. Objectives can be any measurable feature of a model response, including firing rate, spectral power, synchrony, or empirical spectrotemporal similarity to observed neuronal data. In its self-supervised form, GSDR combines a supervised objective term with an activity-dependent MCDP term. The alpha parameter controls the mixture between these update paths, and the genetic component stores the best model state while deselecting states that exceed a loss threshold. We show that GSDR can train both simple and more complex spiking models (Figs. 2-6), including beta/gamma spectral shifts and stimulus-induced gamma dynamics from macaque monkey visual cortex recordings. We show simulations with Izhikevich neurons to test model-class robustness. This shows that GSDR is a methodological framework for automated exploration of nonlinear neural models, with predictive routing serving as a biologically meaningful application.

## Results

### Evolutionary exploration with GSDR

The overall strategy of Genetic Stochastic Delta Rule (GSDR) is to combine objective-guided optimization, stochastic exploration, an activity-dependent update term, and genetic selection/deselection (Fig. 1; see also Materials and Methods, Algorithm 1, Equations 2.1-2.2). The neuronal network model generates a series of neural responses (X[t], membrane potential of all neurons across time on trial t). In the activity-dependent path, these voltages define an MCDP representation R[t], which depends on membrane-potential similarity over time and is constrained by the existing model connectivity. In the supervised path, metric functions are applied to the neuronal response to calculate model-derived metrics (M[t]:= F(X[t])). The distance between each metric and its target value (T[t]) defines the loss or reward value for the current trial (L[t]:= Eval(M[t], T[t])). The trainable model parameters U[t] are then updated using a mixture of the supervised term and the activity-dependent term. The alpha parameter controls this mixture: alpha = 0 corresponds to a fully supervised update, whereas alpha = 1 corresponds to an update driven by the activity-dependent MCDP term and stochastic exploration. In dynamic-alpha simulations, alpha follows a bounded stochastic random walk, but it is still filtered by the same selection/deselection rule as other parameters. The genetic component stores the best model state found so far and reverts to that state when the current loss exceeds the threshold.

**Fig 1:**
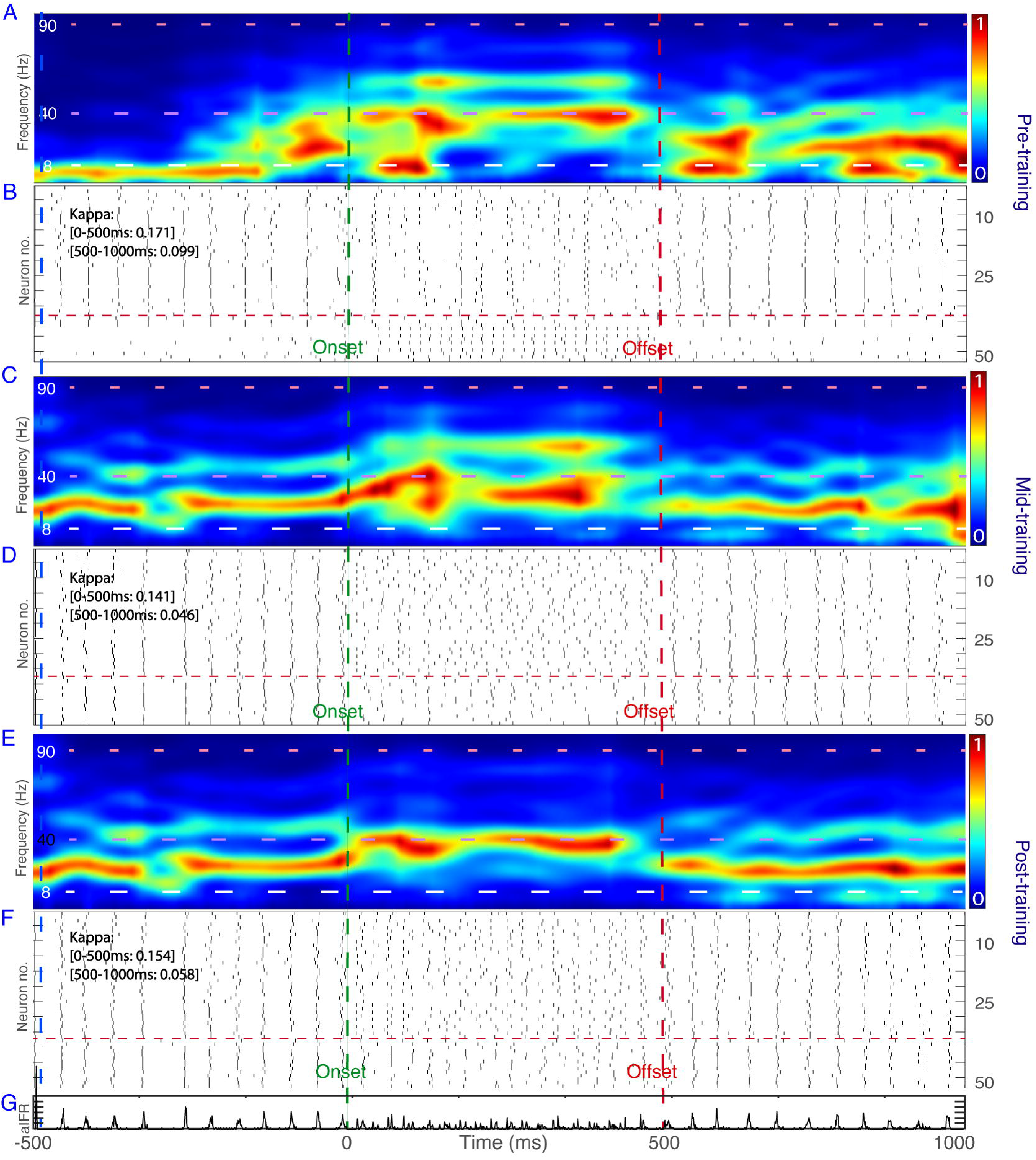
Flowchart of optimization with GSDR. At each iteration, a simulation is performed to obtain neural signals, here membrane potentials over time. A metric or set of metrics is measured from these signals, such as power spectral density, firing rate, or synchrony. The distance between measured metrics and target metrics defines the current loss and contributes to the supervised update path. In parallel, MCDP computes an activity-dependent representation from pairwise membrane-potential similarity. The model updates parameters by mixing the supervised and activity-dependent terms according to alpha. Alpha can be static or dynamic. In the dynamic implementation, alpha follows a bounded stochastic random walk and is subject to the same selection/deselection logic as other parameters. At any step, if the ratio between current loss and selected optimal loss exceeds a threshold, the next trial resets to the selected model state.

**Fig 2:**
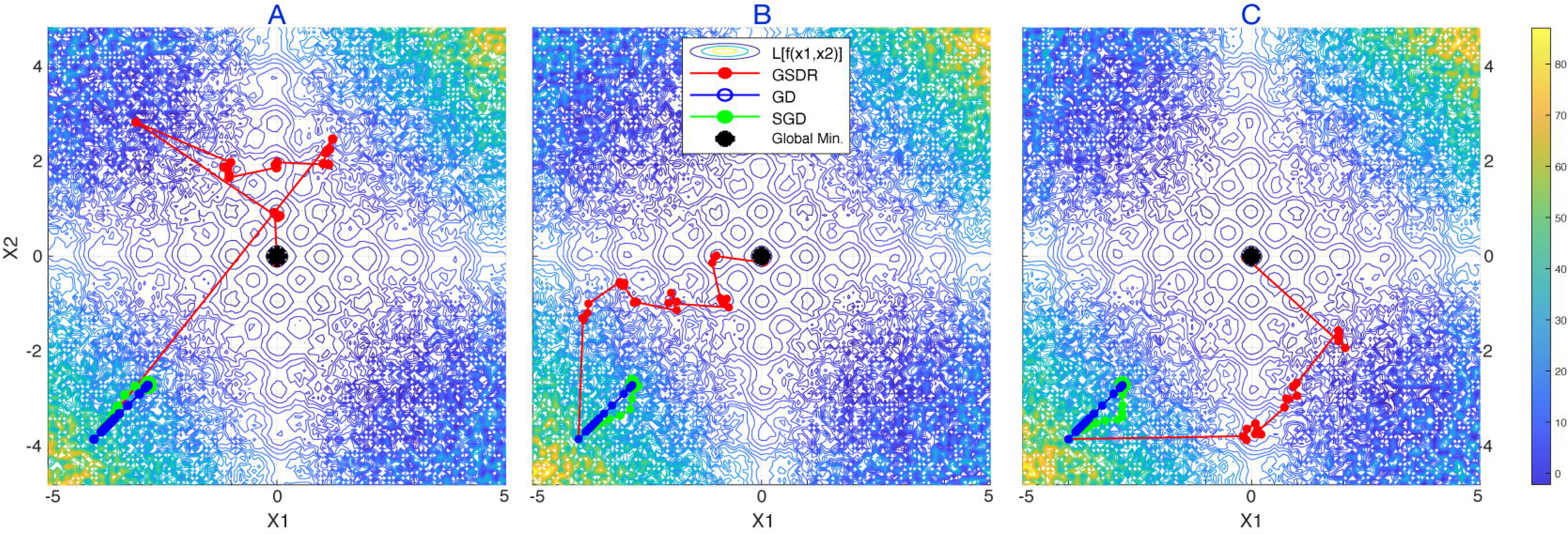
Optimization in a noisy Rastrigin function. Three separate runs (A,B,C) are shown for 1000 iterations for each optimization method, with their corresponding progress. Notice the advantage of gradient-free exploration (red, GSDR) versus gradient-based methods (blue, Gradient Descent - GD, green, Stochastic Gradient Descent - SGD). Stochastic exploration (GSDR) functioned well despite a parameter-loss space that is highly non-convex and noisy. Gradient - based methods tended to get stuck in local minima.

### Advantage of stochastic exploration in noisy, non-linear spaces

We first show the advantage of evolutionary strategies in a modified version of the Rastrigin function [77,78]. This simulation illustrates the stochastic and genetic search behavior of GSDR in a low-dimensional setting before applying the framework to neural circuit models. Since neurobiological circuits contain stochastic and nonlinear elements, we modified the Rastrigin function by adding a uniform noise process. We simulated a 1000-iteration run on the noisy Rastrigin function for GSDR, Gradient Descent (GD), and Stochastic Gradient Descent (SGD). The objective of each optimization function was to find the global minimum. The results (Fig. 2) indicate that GSDR, as an evolutionary algorithm, can be effective in non-convex and noisy parameter-loss spaces, similar to other methods from the same family [69,76–80]. Although this is a simple two-parameter search space, it provides an accessible demonstration of the search logic before the later spectral and empirical neural-model applications.

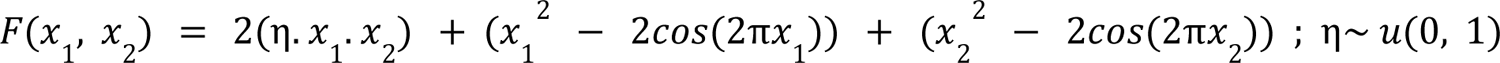

### Pedagogical spiking-network demonstrations

These examples show how GSDR behaves in progressively more neural settings, including single-neuron membrane-potential and firing-rate tuning (Supplemental Fig. 1-5).

### Spectral push-pull through synaptic self-modulation

We next simulated an oscillatory push-pull interaction in an E-I neuronal population. We trained the model to express a flexible switch between beta and gamma spectral states, motivated by the oscillatory motifs described in predictive routing [47,81,82]. In this simulation, the architecture is related to the supplemental E-I models (Supplemental Figs. 4-5), but synaptic connection weights are included as trainable variables to increase the dimensionality of the parameter space. The objective was defined as a spectral ratio under two input conditions. In condition 1, used as a sensory-processing-like condition (Fig. 3C), the objective was for gamma-band power (35-100 Hz) to exceed beta-band power (15-30 Hz). In condition 2 (Fig. 3D), the objective was for beta-band power to exceed gamma-band power.

**Fig 3:**
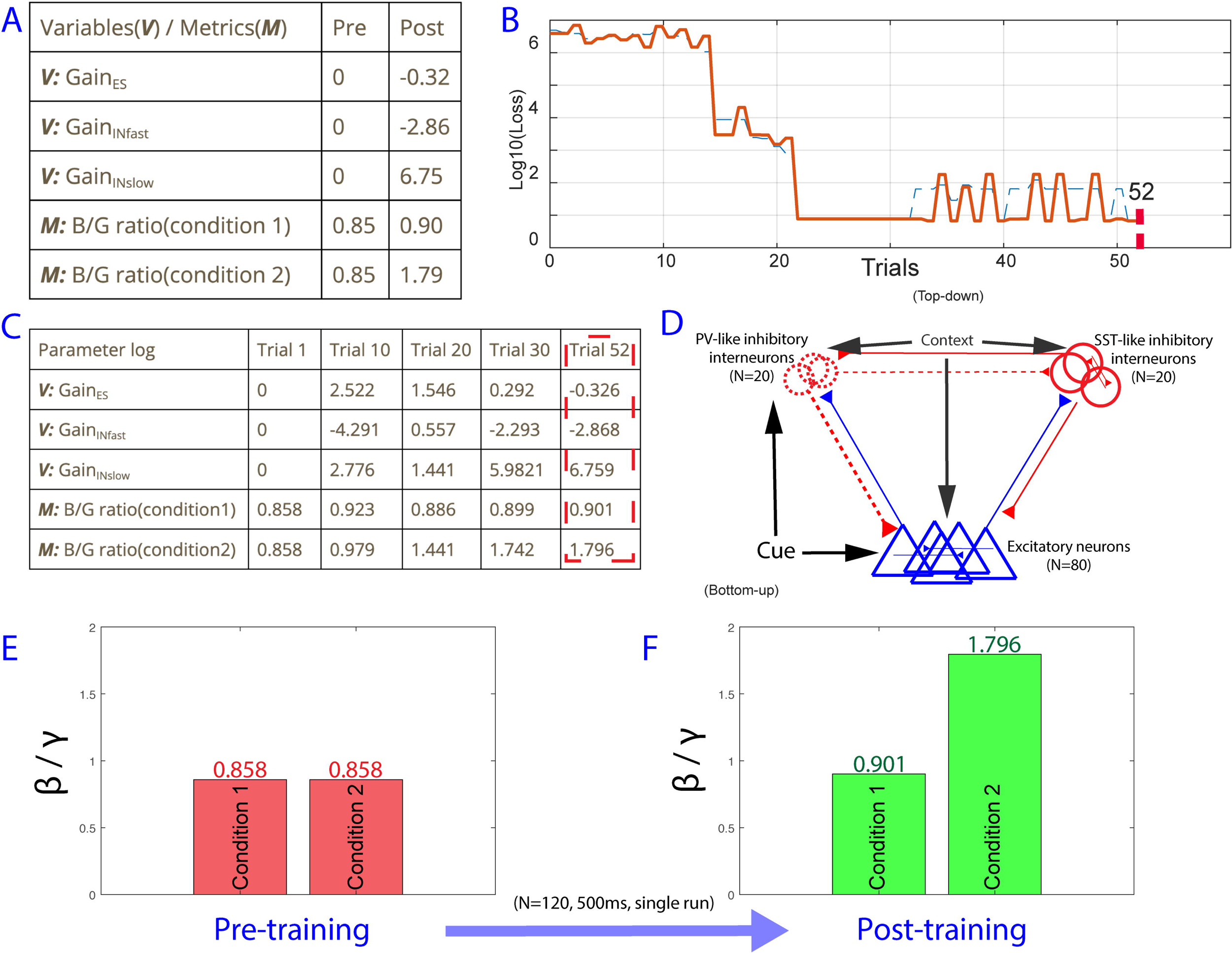
E-I population spectral power ratio tuning. Broad beta [15-30 Hz] average power is compared with broad gamma [40-90 Hz] average power. (A) Variables and metrics before, during, and after training. (B) Network schematic. (C) Post-training raster plot in condition 1. (D) Post-training raster plot in condition 2.

The model achieved the conditional objective (Fig. 3): gamma-band power exceeded beta-band power in condition 1, whereas beta-band power exceeded gamma-band power in condition 2. The total synaptic gain, defined as the sum of all synaptic connectivity weights, changed minimally, whereas the variance and distribution of synaptic weights changed to match the objective. This result supports a constrained sufficiency claim: within the synaptic-weight hypothesis space, recurrent synaptic plasticity can modulate the relative spectral response of a neural circuit without changing synaptic time constants or population-average gain.

In addition, the transition from gamma (condition 1, Fig. 3C) to beta (condition 2, Fig. 3D) resulted in substantially suppressed population activity, especially among excitatory cells. We interpret this as a spectral/circuit-level phenotype that is consistent with prior proposals linking beta states to reduced spiking activity [83] and with predictive routing models [47,48].

### Contextual spectral push-pull with top-down modulation

Predictive routing proposes that top-down inputs carrying predictions through beta-band activity can suppress or modulate bottom-up gamma-band processing [47]. We therefore modeled how top-down inputs can modulate a lower-order network state from a gamma-dominated to a beta-dominated spectral regime. We did not model two cortical regions simultaneously; instead, we modeled top-down input to a hypothetical lower-order area, similar to previous work on top-down effects of attention on sensory processing [42]. In this simulation, the model’s objective was to tune the relative beta/gamma power ratio (Fig. 4). This task is related to the previous spectral-ratio model (Fig. 3), but local synaptic weights were fixed. Instead, top-down contextual gain parameters were allowed to change, as hypothesized in predictive routing. These gains modulated the strength of top-down synaptic input to excitatory cells, slow inhibitory interneurons (SST-like), and fast inhibitory interneurons (PV-like) (Fig. 4B).

**Fig 4:**
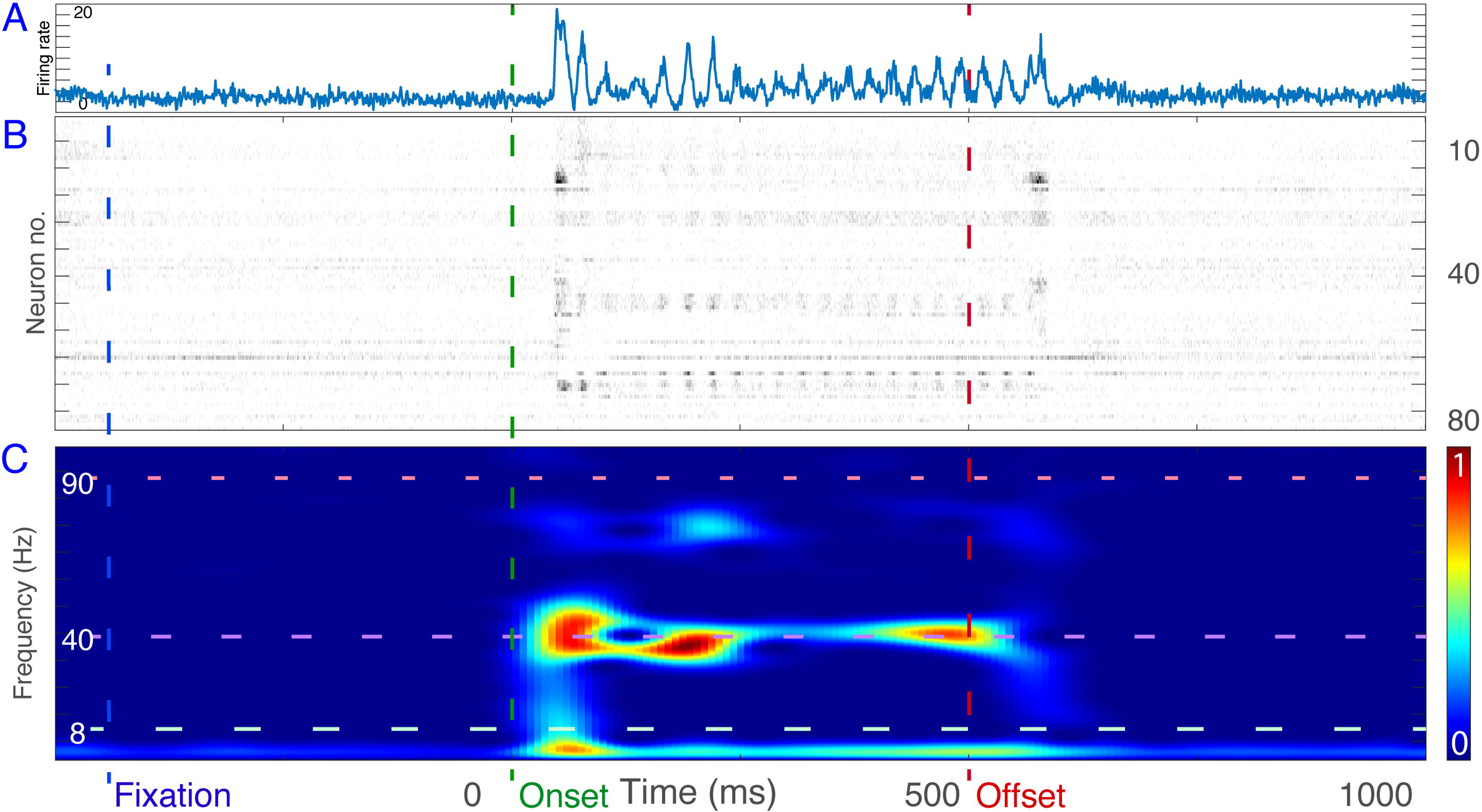
E-I contextual spectral relative power tuning. Beta [∼10-30 Hz] average power is compared with middle gamma [55-75 Hz] average power. (A) Summary of pre- and post-training variables and metrics. (B) Loss across trials relative to target metrics. (C) Summary of training variables and metrics during training. (D) Network schematic. (E) Spectral response before training. (F) Raster plot before training. Due to stochasticity from internal noise and receptor-synaptic interactions, the beta/gamma ratio in condition 1 varies slightly between pre- and post-training simulations.

Top-down input gain to all populations was initially set to 0 for both conditions. In condition 1, top-down inputs remained 0, such that the model stayed near the baseline gamma-dominated state, operationalized as a beta/gamma ratio below 1 (Fig. 4E). Next, GSDR was allowed to modulate top-down gain parameters with the objective of shifting the model toward a beta-dominated state in condition 2. GSDR found a solution with positive gain on the slower inhibitory population and negative gain for the fast inhibitory population (Fig. 4C, red box). Post-training responses showed a context-dependent spectral power shift (Supplemental Fig. 3C-F), with respect to the beta (∼10-30 Hz) to gamma (>35 Hz) power-ratio objective, ending at a ratio of ∼1.8 (Fig. 4F). These gain parameters are consistent with studies suggesting that top-down feedback can drive slow inhibitory interneurons [18,84] and shift networks toward beta-dominated states [47,85].

### Narrow-band gamma oscillations are observed in electrophysiology recordings from the primate visual cortex

The previous contextual model (Fig. 4) generated overly synchronous oscillations (Supplemental Fig. 3D) that do not resemble typical in vivo electrophysiology. To address this, we used synaptic plasticity to model electrophysiologically recorded, visually induced gamma oscillations (∼38 Hz) from the visual cortex of awake macaques. The objective was to let GSDR modulate synaptic connectivity to reproduce observed gamma-band neural dynamics across stimulus-off and stimulus-on periods. Current models of oscillatory dynamics, including Pyramidal-Interneuron Network Gamma (PING), can explain gamma-band dynamics and distinguish between strong and weak regimes [42,58–60]. In strong PING, gamma-band oscillations (>35 Hz) require strong input drive and feedback inhibition with fast inhibitory time constants, yielding highly synchronous rhythms that dominate individual-neuron responses. In weak PING models [58–60], noise or heterogeneous synaptic weights generate population-level gamma with sparse and irregular firing in pyramidal cells.

Prior to the simulations, we examined whether real neurons display a strong or weak PING phenotype in vivo. We performed electrophysiological recordings in the visual cortex of awake macaque monkeys during presentation of drifting grating stimuli, known to induce gamma oscillations (see Methods for details). We used neuronal data from area MT/MST because this area was highly sensitive to moving grating stimuli, and we consistently observed a strong narrow-band gamma response during stimulus presentation (Fig. 5). The peak frequency of this visually evoked oscillation increased over the session by about 2.5 Hz (Supplemental Fig. 14), which replicates earlier work showing an increase in gamma peak frequency from early to later trials in a session [135]. In the same session, we also recorded from area PFC with the same type of recording electrode (upper sub-panel, Supplemental Fig. 14). PFC did not show gamma oscillations and instead had a dominant peak frequency in the beta band (∼20 Hz), consistent with prior electrophysiology showing beta as a dominant frequency of higher-order cortex [83].

**Fig 5:**
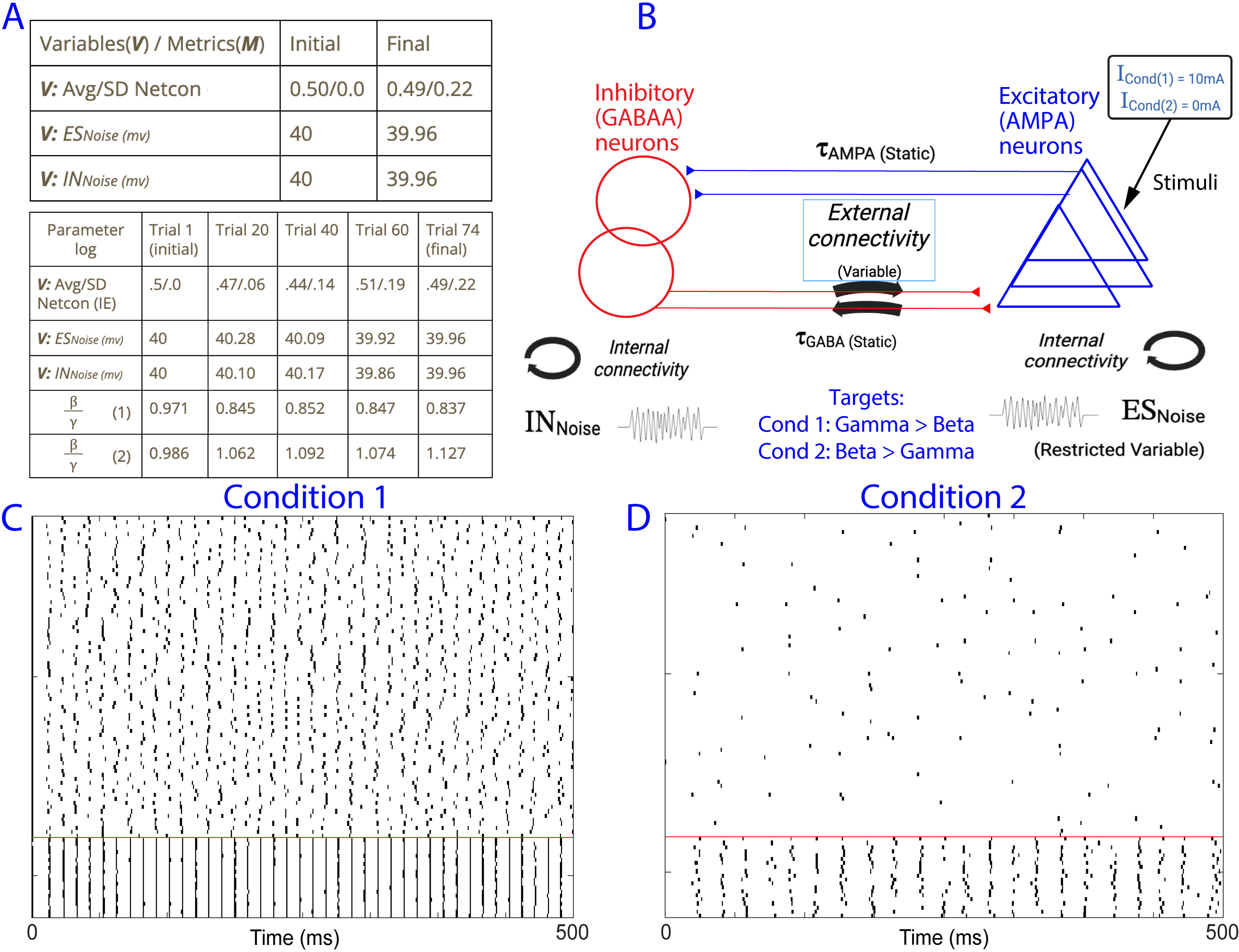
Single-unit neural response from visual areas MT/MST (NHP). (A) PSTH and all-trials cumulative raster plot (100 trials, 80 neurons). (B) Trial-average cumulative raster plot of neurons. Blue line: fixation; green line: stimulus onset; red line: stimulus offset. (C) Corresponding time-frequency response, scaled from 0 as the lowest power to 1 as the highest power across the duration shown above, showing a stimulus-dependent evoked spectral response around ∼37.5 Hz.

**Fig 6:**
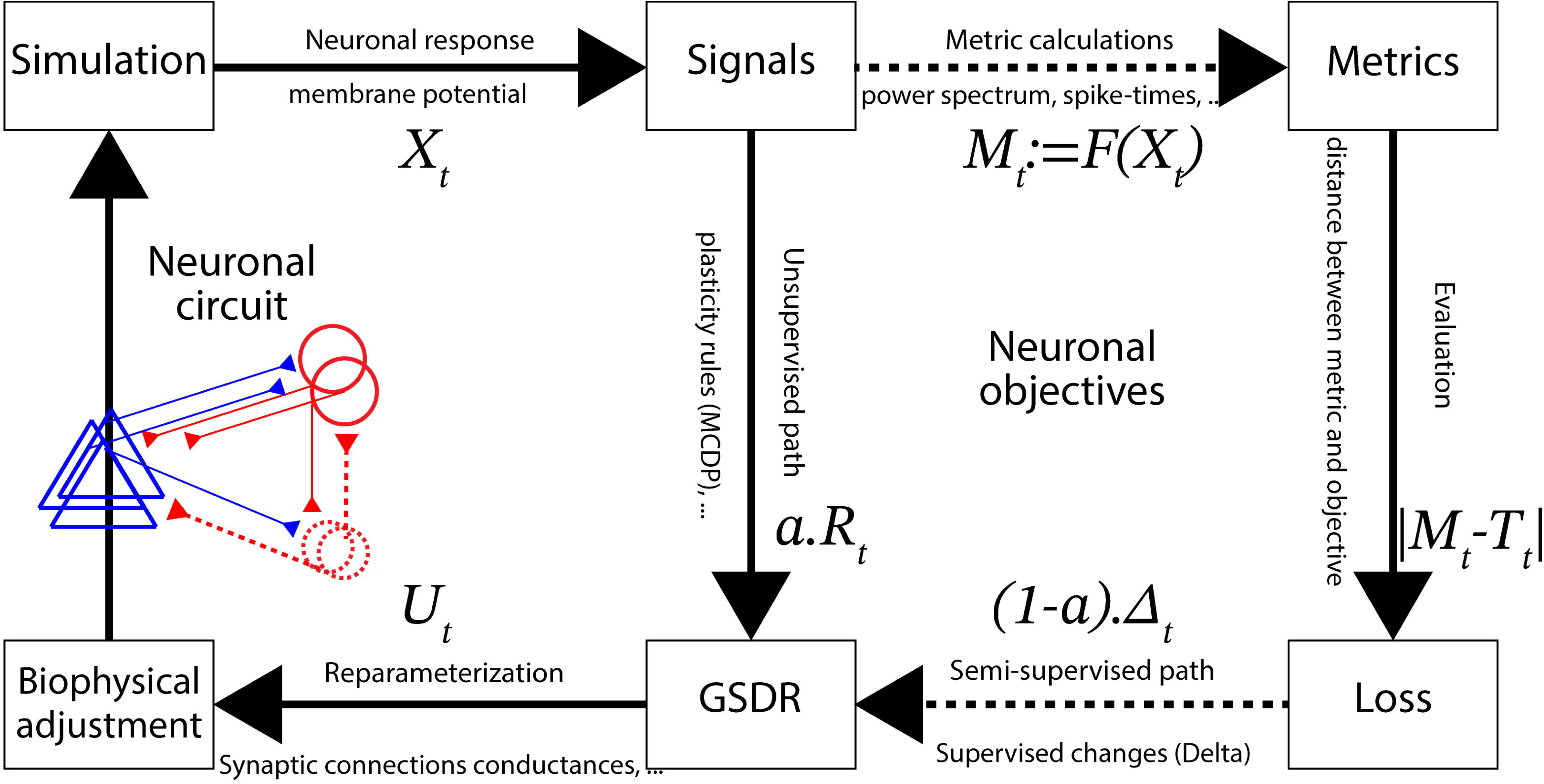
Single-unit neural responses from the model. (A) Time-frequency response of the model before training, scaled from 0 as the lowest power to 1 as the highest power and 1/f adjusted. (B) Corresponding raster plot before training, reflecting highly synchronized spiking. (C) Time-frequency response during mid-training (trial ∼200). (D) Corresponding raster plot during mid-training. (E) Time-frequency response after training. (F) Corresponding raster plot after training. (G) PSTH of all neurons in panel F. During the stimulus-on condition, stimulation was randomly provided to half of the excitatory neurons from 0 ms (onset, green dashed line) to 500 ms (offset, red dashed line). Network synchrony based on the kappa value [136] is also noted for each raster, showing less synchrony than the strong-PING control (Supplemental Fig. 15) and reduced kappa during stimulus presentation.

### Self-supervision guides parameter-space exploration

To study circuit mechanisms that can produce this gamma response, we implemented a model with 50 neurons (36 excitatory, 14 inhibitory). Instead of relying on tunable synaptic time constants, inhibitory neurons were defined by two connectivity classes [21,86]. One class had locally focused connectivity (PV-like, restricted to local synaptic connections with ∼10% of total neurons; Supplemental Fig. 6), whereas the other class had global widespread connectivity (SST-like, with no restriction on the number of synapses to other neurons). Unlike classic PING models that rely on fast inhibitory time constants for gamma, we did not modulate synaptic time constants in this model. AMPA and GABA rise and decay constants were fixed [87,88]. Instead, GSDR was allowed to modulate recurrent synaptic conductances. This was a hypothesis-driven choice: recurrent E/I weights are a plausible circuit substrate through which local networks can alter spectral responses without changing membrane-channel kinetics or synaptic time constants. The objective was to approximate the spectrotemporal features present in the observed neurophysiology by maximizing similarity between model and data power spectra (Fig. 5A and Supplemental Fig. 12) in both the pre-stimulus and stimulus periods. We refer to this as the spectral similarity task.

For this task, we used the self-supervision component of GSDR (Algorithm 1). The spectrotemporal response of both the model and the data was quantified by taking each neuron’s power spectrum and averaging spectra across neurons (see Methods). The model optimized its spectral response to reproduce the stimulus-evoked gamma as well as the pre-stimulus period spectral response. The loss was calculated using the log-ratio between the model and empirical spectral responses after normalization (Supplemental Fig. 12), along with a one-sided mean exponential error (MXE; Equation 5) penalizing models with very low spike counts. To simulate the visual stimulus, we modeled stimulus onset at time 0 as an increase in external input: a noisy direct-current step function with amplitude 1 nA, convolved with a 120 Hz pulse train and lasting 500 ms. Background noise was modeled as a stationary random current process [89], independently present for every neuron.

Model and neurophysiology data both showed a beta spectral peak (∼15-20 Hz; Supplemental Fig. 7A; see also Supplemental Fig. 12) during the pre-stimulus baseline and a stimulus-induced shift toward gamma (∼35-40 Hz; Supplemental Fig. 7B). The empirical and model spectra early and late in training are shown in Supplemental Fig. 12 for pre-stimulus and stimulus periods. During this training, recurrent synaptic conductances were the changing parameters; during the simulated stimulus window, those conductances were fixed and the beta-to-gamma transition was driven by stimulus input interacting with the learned circuit.

As an additional model-class robustness test, Izhikevich simulations reproduced corresponding baseline, stimulus-on, and PSD-level spectral behavior (Supplemental Fig. S16, baseline; Supplemental Fig. S17, stimulus-on; Supplemental Fig. S18, PSDs), showing that GSDR is not limited to the conductance-based implementation.

These results show that GSDR can optimize spiking models for the spectral similarity task without relying solely on external supervision or manual parameter dictation. Recapitulating the observed gamma-band response did not require manual adjustment of synaptic time constants or strong uniform inhibitory-to-excitatory connections. Instead, the post-training state was reached through sparse and non-uniform recurrent synaptic weights (Supplemental Fig. 6A-B). This supports the conclusion that GSDR can search a constrained synaptic-weight space and identify candidate circuit configurations with weak-PING-like properties: the oscillation is most prominent at the population level rather than dominating every single neuron (Figs. 5-6).

## Discussion

Our work establishes GSDR as an evolutionary optimization framework for nonlinear spiking and neural models. GSDR combines a supervised objective term, stochastic exploration, an activity-dependent MCDP term, and a genetic selection/deselection mechanism that retains selected model states and rejects states exceeding a loss threshold. The main contribution is methodological. Predictive routing provides a biologically meaningful case study because it involves oscillations, spiking activity, and beta/gamma state changes. We used GSDR in three main applications: tuning recurrent E-I connectivity to modulate beta/gamma spectral ratios (Fig. 3), tuning top-down contextual input gains to shift the same network toward a beta-dominated regime (Fig. 4), and fitting model spectra to empirical macaque MT/MST stimulus-evoked gamma dynamics (Figs. 5-6). We also simulated Izhikevich networks (Supplemental Fig. S16, baseline; Supplemental Fig. S17, stimulus-on; Supplemental Fig. S18, PSDs), showing that the framework is not tied to the conductance-based model class.

### What has GSDR and evolutionary approaches taught us about biophysical models?

Many nonlinear neural models perform computations [90–92], but their parameter spaces often require extensive and complex search. Self-supervised learning, stochastic exploration, and genetic selection/deselection is an effective strategy to search this space. The primary novelty of GSDR is not that E/I networks can generate oscillations, which is well established, but that GSDR automates the search for parameter regimes that satisfy spectral and empirical objectives under explicit constraints. The alpha parameter controls the mixture between the supervised objective term and the activity-dependent MCDP term. In the dynamic-alpha implementation, alpha is not optimized by a separate objective; it follows a bounded stochastic random walk and is retained or rejected through the same selection/deselection logic as other model quantities. The resulting connectivity patterns (Supplemental Fig. 6B) were found within a hypothesis space and were not presented as an unconstrained discovery of all possible biological mechanisms. Parameter bounds, such as non-negative conductances and maximum synaptic strengths, were used to exclude physically or biologically invalid regimes rather than to force a particular spectral solution.

Previous studies [42,55,56,59,92] have shown that E/I networks can generate gamma and beta rhythms, including strong and weak PING-like regimes [53,55,58–61]. Our novelty is therefore not the existence of such rhythms. Instead, the present work shows that a hybrid evolutionary optimizer can search for bounded parameter regimes that reproduce spectral objectives and empirical spectrotemporal targets. Our empirical MT/MST recordings were more consistent with sparse, irregular, population-level gamma than with a globally synchronous strong-PING phenotype (Fig. 5). In the spectral similarity task, the post-training model also exhibited sparse participation across individual neurons (Fig. 6), with narrow-band gamma during stimulation and beta-dominated activity during baseline (Supplemental Fig. 12). The added Izhikevich simulations further show that similar qualitative spectral behavior can be evaluated in the Izhikevich model (Supplemental Figs. S16-S18).

### What has GSDR taught us about predictive routing and oscillatory shifts?

Our results showed that top-down contextual gains can be optimized to shift a network from a gamma-dominant to a beta-dominant spectral state (Fig. 4). This result is consistent with predictive-routing ideas in which top-down, preparatory signals modulate sensory processing, but it should not be interpreted as a complete predictive-processing computation. The model does not perform explicit mismatch detection, belief updating, omission detection, or behavioral prediction. A formal task-performing predictive-routing model remains a future application of the framework.

We also tested whether local recurrent synaptic weights are sufficient to induce beta/gamma spectral shifts under fixed cellular and synaptic-kinetic assumptions (Fig. 3). This was a hypothesis-driven search, not an unconstrained search over all possible mechanisms. Alternative implementations could include changes in intrinsic excitability, synaptic kinetics, neuromodulatory gain, or targeted external input. The present result therefore shows that GSDR can identify bounded parameter regimes in which the same network expresses different spectral regimes under different input conditions.

### Connections to other computational modeling frameworks

Several computational frameworks can be used to fit observed neural dynamics, including Dynamic Causal Modeling (DCM) [98–101], neural mass models [19], and conductance-based spiking networks. These approaches occupy different points on a modeling spectrum. Neural mass models provide interpretable population-level dynamics and may reproduce many spectral phenotypes with fewer state variables. Reduced-spiking models provide cell-level dynamics while preserving spike timing and E/I interactions. Conductance-based models provide access to channel-, receptor-, and current-level hypotheses.

When performing computational modeling, it is important to select a model with sufficient detail to capture the desired behavior. This does not always require the level ionic-level mechanisms in the Hodgkin-Huxley equations. We present GSDR as a model-class-flexible optimization framework. The Hodgkin-Huxley-style implementation is one compatible model class, useful when conductance-level hypotheses are central. The Izhikevich simulations provide a middle ground, showing that GSDR can also operate within simpler models while preserving spike timing and E/I interactions (Supplemental Figs. S16-S18). Future neural mass implementations would test the same optimization logic at a population scale. More exhaustive benchmarking across model classes and optimizers remains an important next step.

### Future extensions of the evolutionary modeling framework

Evolutionary strategies open an alternative class of algorithms for computational modeling in neuroscience. GSDR can be used to compare how different model classes and objective functions reproduce neuronal data, including objectives derived from predictive coding, predictive routing, or other theories. The present work focuses on oscillatory and spectrotemporal objectives because these are central to the predictive-routing case study, but the framework is not limited to that theory. Future work can apply GSDR to explicit task-performing models where the functional computation can be tested directly rather than inferred from spectral correlates.

In principle, any observed neurophysiology feature that can be measured from a model or data can be used as a GSDR objective. This includes LFP-spiking relations, spike timing and rate coding, spike-field coherence (SFC) [110], neuronal metabolism and neuromodulator interactions [23,105,111], multiple co-existing gamma oscillations [112–114], and distinct mechanisms for beta versus alpha oscillations [115–118]. The models in the present manuscript contain relatively few neurons (∼100 neurons), although smaller models can be augmented to simulate larger dynamics (for a 1,000-neuron augmentation, see Supplemental Fig. 13). Future work should extend these models to multiple areas and larger numbers of neurons and trainable parameters [118], especially as modern neurophysiology enables thousands of neurons to be recorded during task performance [119].

## Conclusion

To summarize, GSDR can search neural parameter spaces, reduce reliance on manual tuning, and reproduce spectral objectives using multiple neuronal model types. We propose that GSDR be applied to different objectives and datasets in an iterative model-data virtuous cycle.

## Materials and Methods

Here we used the DynaSim toolbox (Matlab) and Jaxley toolbox (Python) to implement neural circuit models [92,127]. These models consist of one or multiple networks of neurons with different cell types. The main simulations use conductance-based spiking neurons with synaptic mechanisms, and the revision adds Izhikevich simulations as a model-class robustness analysis. Additional mechanisms can be included when needed, such as receptor, ion-channel, synaptic, or input-current terms. The system of equations governing membrane-potential dynamics is written in Equation 1

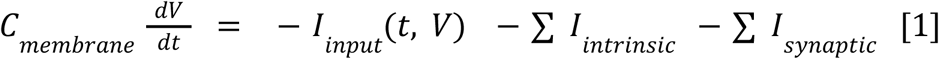

*Here t is time, V_i is the membrane potential of neuron i, C_m is membrane capacitance, I_int denotes intrinsic membrane currents such as sodium, potassium, and leak currents, I_syn denotes synaptic currents from other neurons, and I_input denotes external or stimulus-related input current. The sign convention is that positive outward current hyperpolarizes the membrane in this formulation. This framework allows models to include multiple populations, cell classes, and synaptic mechanisms such as AMPA, NMDA, and GABA currents*.

Genetic Stochastic Delta Rule (GSDR), inspired by the stochastic delta rule [74] and genetic optimization [69], explores trainable model parameter spaces and deselects suboptimal states to achieve one or more objectives. Objectives are target values for metrics derived from the model, such as membrane potential, firing rate, synchrony, or spectral population dynamics. The general update is summarized in Equations 2.1 and 2.2.

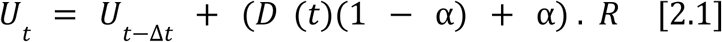

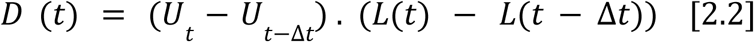

Here U_t represents the trainable model parameters at trial t, delta(lambda) is a stochastic exploration term controlled by the exploration factor lambda, L_t is the current loss, D_t is the temporal difference used by the supervised update logic, alpha_t is the mixing factor between supervised and activity-dependent update paths, and R_t is the MCDP activity-dependent representation. Alpha ranges from 0 to 1. When alpha = 0, the update is fully supervised by the objective-dependent term. When alpha = 1, the update is driven by the MCDP activity-dependent term and stochastic exploration. In static-alpha simulations alpha is fixed. In dynamic-alpha simulations alpha follows a bounded stochastic random walk and is retained or reverted through the same genetic selection/deselection logic as other trainable quantities. We used stochastic delta-rule updates and, for comparison, SGD-based updates implemented in JAX [128,130].

**Alg1:**
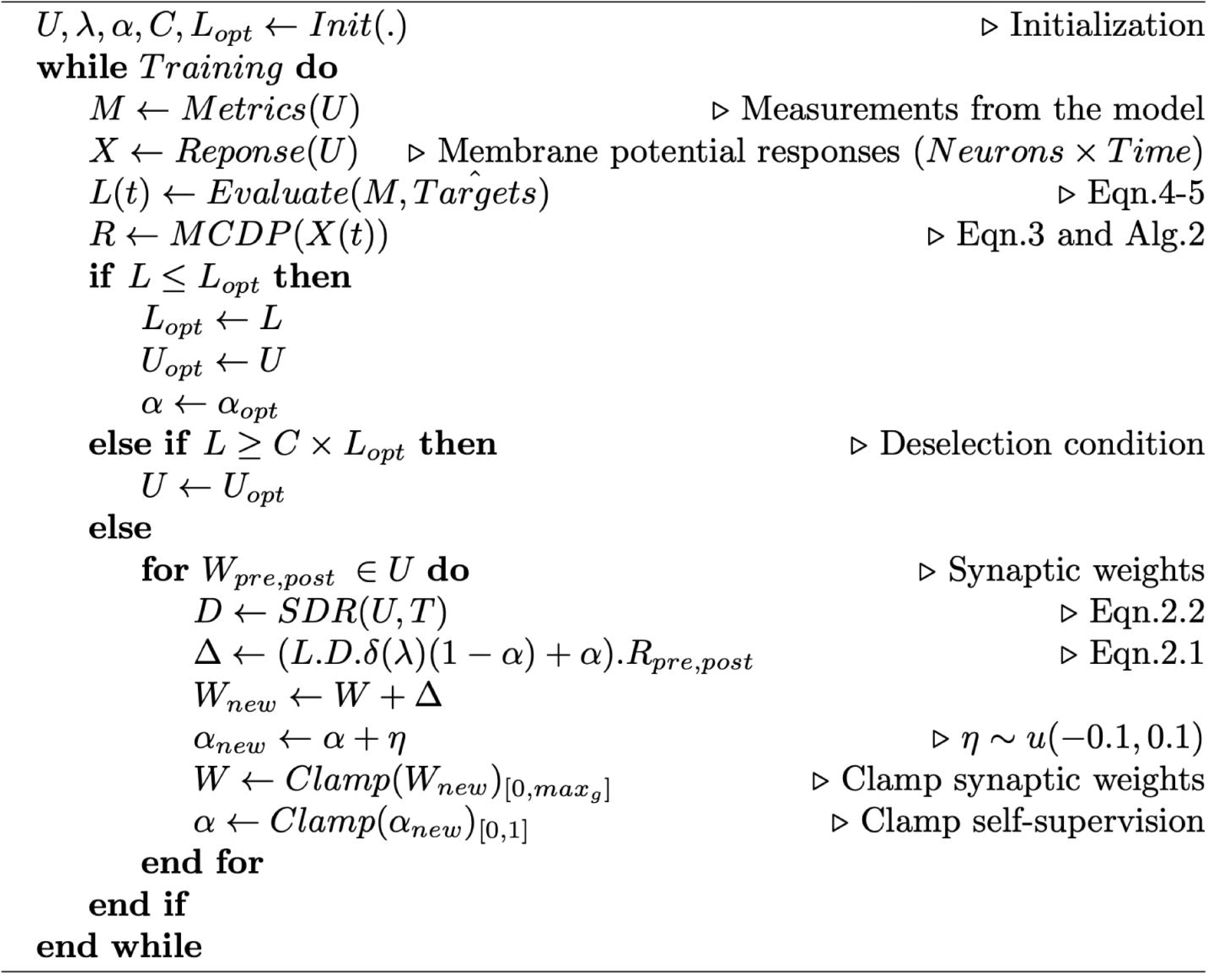
GSDR pseudocode. Trainable model parameters are included in the U. After initializing model and optimization parameters (alpha: self-supervision factor; lambda: exploration factor; C: deselection threshold), the training loop begins. At each trial, simulations are performed with the current parameters while recording membrane potentials X. Metrics are calculated from X to define the loss, and activity-dependent MCDP factors R are calculated from X where applicable. If the batch loss is better than the previously selected loss, the parameters are stored as the selected model state. If the batch loss exceeds C times the selected loss, the model reverts to the selected state. For each trainable parameter set in U, the update step is applied (Equations 2.1-2.2). When MCDP is not defined for a parameter, R is set to 1. Alpha can also change by bounded stochastic steps and is retained or reverted through the same selection/deselection rule. Parameters are clamped to valid ranges, such as non-negative synaptic conductances and maximum allowed strengths, to prevent physically or biologically invalid values rather than to force a specific solution.

MCDP takes membrane-potential responses X as input and calculates the correlation between pre-synaptic potentials and time-shifted post-synaptic potentials. The post-synaptic traces are shifted according to the corresponding synaptic time constant, such as tau_AMPA or tau_GABA, to account for synaptic timing. The output is an N by M matrix for N pre-synaptic and M post-synaptic neurons. This matrix is an activity-dependent representation constrained by the existing model connectivity. It is related to STDP in that it uses temporal relationships between pre- and post-synaptic activity, but it is computed from membrane-potential similarity rather than discrete spike times alone. The calculation is summarized in Equation 3.

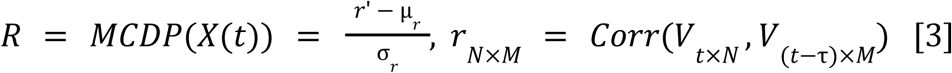

The value of R is set to 1 for any trainable variable for which MCDP is not defined, including parameters that do not correspond to a pre-post synaptic pair.

**Alg2:**
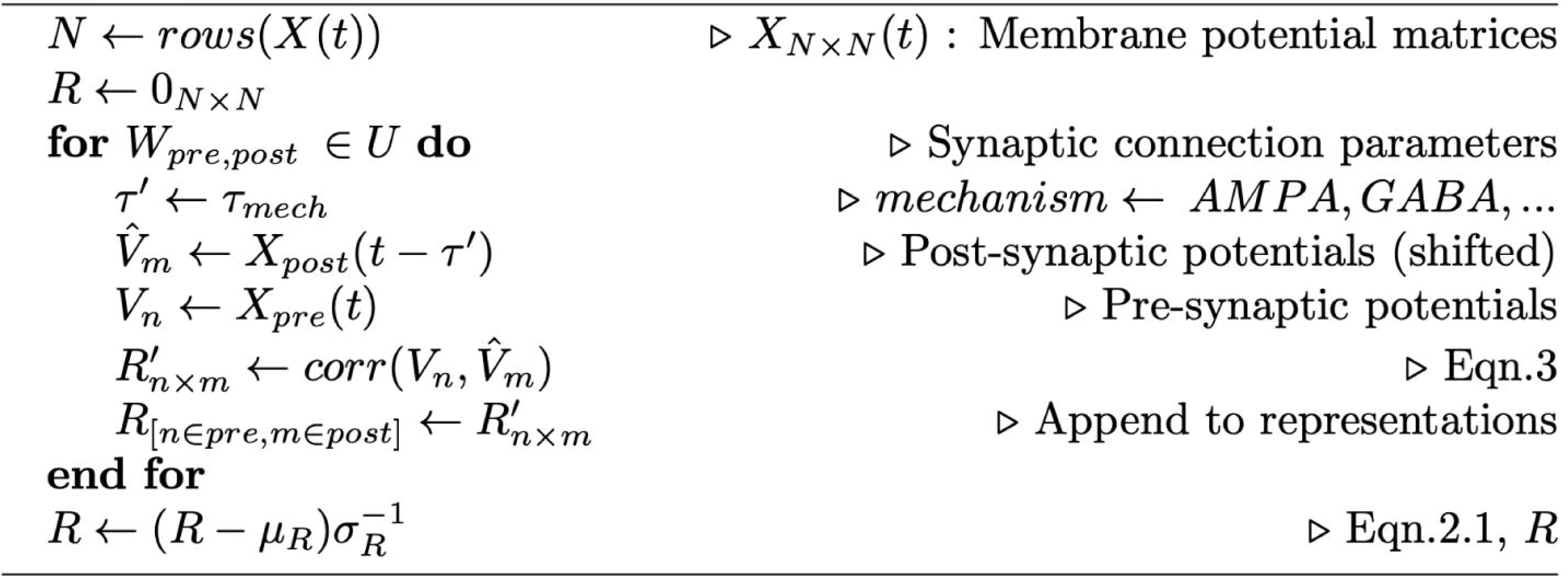
MCDP pseudocode. For trainable parameters shared between a pre-synaptic and post-synaptic neuron, the membrane-potential matrix X is used to calculate cross-correlation. For the post-synaptic matrix, signals are shifted according to the relevant synaptic time constant to account for timing relative to the pre-synaptic reference signal. After all MCDP factors are calculated, the normalization step (Equation 3) is performed and the activity-dependent representation matrix R is returned for use in the GSDR update (*Algorithm 1*, Equation 2.1).

GSDR can include objectives, such as specific firing rates, spectral or temporal patterns. This requires defining the corresponding metric-target for each objective. A metric (*M*) can be a direct or indirect measurement from the model, a function that is applied on the model’s response. A metric generally consists of three arguments; spatial annotation (which part of the circuit to be measured), temporal annotation (when in time points or intervals of simulation should be selected for this measurement) and the measurement function (what aspect of the annotated model output should be measured). In our simulations we have utilized both metrics that have a biophysical unit (current and voltage) as well as metrics that are derived from the model’s response (firing rate and spectral response). Algorithm 1 describes how GSDR uses these metrics to train models.

*Objectives consist of one or more target values (T). The loss function in the single-neuron simulations (Supplemental Fig. 1) and the pedagogical population simulations (Supplemental Figs. 4-5) was calculated by mean square error (MSE) between the measured metric and the target. In the population firing-rate tuning simulation (Supplemental Fig. 4), two objective components were used: MSE between the excitatory population firing rate and its target, and MSE between the inhibitory population firing rate and its target. For population natural-frequency tuning (Supplemental Fig. 5), the loss was calculated by MSE between the measured natural frequency, defined as the peak-power frequency, and the assigned target. In the population spectral power-ratio task (Fig. 3) and contextual top-down spectral task (Fig. 4), the objective was defined on the beta/gamma power ratio*.

In the spectral similarity simulations comparing the model response to electrophysiology (Figs. 5-6; Supplemental Fig. 12), we defined a loss function by calculating the log ratio of the normalized model spectral response to the normalized target spectral response (Equation 4). Spectral analyses involving power spectral density (PSD) of model and electrophysiology data were performed on continuous neural traces and then averaged across neurons at each frequency bin.

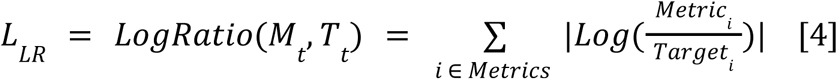

In addition, we defined a one-sided loss function (eqn. 5, mean exponential error, MXE) penalizing models with a very low spike count. This was necessary because without this penalty, some models produced little to no spiking.

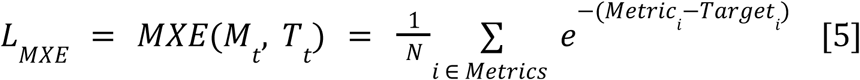

In neural models, it is important to consider the plausibility of objectives and parameter bounds. Some target states can violate biological assumptions, such as nonphysiological membrane potentials, negative conductances, or unrealistically high firing rates. In such cases, the model may not be able to achieve the target within valid parameter ranges. Boundary conditions were therefore used to keep parameters within a plausible and stable subspace. For example, synaptic conductances were constrained to be non-negative and scaled to a maximum strength of 1 (a.u.). These bounds were used to exclude invalid regimes rather than to predetermine a specific spectral solution.

### Integrating GSDR into neural network models

As a basic step, we use a spiking neural circuit model to generate membrane-potential dynamics over time. For single-neuron parameter optimization tasks (Supplemental Fig. 1), GSDR reduces to the supervised stochastic delta-rule case because only the supervised path is included. For population-level tasks, GSDR can use both supervised and activity-dependent paths (Fig. 1). In the activity-dependent path, membrane potentials define synaptic co-activation patterns through MCDP. In the supervised path, model-derived metrics are compared with prespecified objectives. The difference between model performance and the objective defines the loss value for the current trial. GSDR combines this loss-guided update with MCDP, stochastic exploration, and genetic selection/deselection to update model parameters over repeated trials. Importantly, GSDR changes parameters across optimization trials; once the model state is selected, the within-trial spectral transition is driven by the input condition interacting with the fixed learned circuit, not by rapid within-trial synaptic rewiring.

We included basic GSDR examples in the Supplement to illustrate the overall logic of the method (Supplemental Figs. 1-5; Fig. 1). In the first single-neuron example, the model was trained to modulate membrane potential from an initial value of −64.9 mV to a target of −60.0 mV (Supplemental Fig. 1A). The trainable parameter was the leak reversal potential (E-leak). The model converged toward the target, and the loss decreased despite transient divergence (Supplemental Fig. 1B). In the second single-neuron simulation, the model was trained to modulate firing rate from 0 spikes/sec to a target of 70 spikes/sec. The model was allowed to tune the leak reversal potential and an external constant current drive. The firing rate approached the target after exploratory behavior (Supplemental Fig. 1E-F).

We also included population simulations as supplemental pedagogical examples. The first used a 100-neuron E-I model (80 excitatory, 20 inhibitory) with AMPA and GABA synaptic mechanisms [87,88], synaptic conductances, synaptic time constants, and intrinsic noise [89]. The model was initialized with uniform connectivity and trained to achieve target population firing rates (Supplemental Fig. 4). A second population example optimized natural frequency using inhibitory synaptic time constants and noise gain (Supplemental Fig. 5). These examples are retained as implementation demonstrations, while the main text focuses on the spectral-ratio, empirical gamma, and Izhikevich robustness results.

Additional robustness simulations: To evaluate whether the GSDR framework was restricted to the original conductance-based implementation, we added Izhikevich simulations [64]. These simulations used cell-class-specific dynamics while preserving recurrent E/I interactions, spike timing, and spectral readouts. The baseline, stimulus-on, and PSD analyses are reported in Supplemental Fig. S16, Supplemental Fig. S17, and Supplemental Fig. S18, respectively. These simulations were not intended to replace the original model or prove a unique circuit mechanism; they serve as a model-class robustness test showing that the optimization logic can be applied beyond the original model class.

### Experimental Design

#### Description of animal use

All experimental procedures were approved by the Vanderbilt University Institutional Animal Care and Use Committee (IACUC). One adult bonnet macaque (Macaca radiata) aged 18 years (7.4kg) was used in this study. All procedures described here were supervised by Vanderbilt’s IACUC to ensure compliance with all local and federal laws and regulations. The monkey was housed in an AAALAC approved facility (Allentown Inc., NJ, Large Animal Primate Housing units) supervised by an AALAS certified laboratory animal technician and staffed by trained caretakers. The animal was cohoused with conspecifics. Within the facility, a comprehensive preventative medicine and veterinary care program is in place that includes routine care, psychological enrichment, and daily observation of animals. Vanderbilt’s facility has been accredited by AAALAC International and is designed to meet the standards and guidelines set forth in the Animal Welfare Act and the Guide for the Care and Use of Laboratory Animals, Eighth Edition.

Enrichment for the animals was provided daily and focused on promoting species-typical behaviors through social housing, complex foraging puzzles, manipulable toys, and, notably, cognitive, touchscreen-based computer tasks. These strategies aim to reduce stereotypic behaviors and improve psychological well-being. Food (macaque biscuits provided twice daily, produce/fruit provided near daily) and water were provided to the animal daily by the laboratory staff and/or the animal care staff.

The monkey was implanted with two 18mm recording chambers (Christ) placed over the visual/temporal and prefrontal cortex and a headpost. The surgery to implant the recording chamber and headpost was performed using sterilized instruments and aseptic technique and included multimodal analgesia (local anesthetic – lidocaine injected subcutaneously before a surgical incision followed by slow-release Buprenorphine and an NSAID for 3 days) and anesthesia (induction with ketamine, followed by maintenance with the gas anesthetic isoflurane). In addition, antibiotics such as cefazolin were given perioperatively. Veterinarians and veterinary technicians monitored the health of the animal daily and communicated directly with each other and with investigative staff regarding the animal’s health at all stages of this research. Monitoring was performed by visual assessment daily and by inspection of detailed records (on feeding patterns, water consumption, and weight) and semi-annually in a comprehensive sedated exam performed by a veterinarian. Altogether, this ensured that there were no signs of pain or distress associated with these experimental procedures (which can be diagnosed by cage-side behavioral changes, changes in weight or changes in food/water consumption) and ensured the animal’s health and well-being were optimized throughout the study. The animal was not euthanized for the purposes of this study.

#### In-vivo neural signal recordings

To insert electrodes into the brain we used a mechanical Microdrive (Narishige, Tokyo, Japan) that was mounted onto the chambers. A guide tube was lowered through a recording grid (Christ) to penetrate the dura mater and granulation tissue. We then acutely introduced a 128-channel linear deep array into area MT/MST. We used a linear 128-channel recording array where the inter-contact spacing was 40 microns (Diagnostic Biochips, Glen Burnie, MD). Recordings were acquired using a RHD System (Intan technologies, CA) sampling at 30 kHz. Recordings were electrically grounded to the guide tube.

Grid positions were pre-determined based on a pre-recording MRI scan with the chambers and grid in place, with water-saline to mark the trajectories of each grid hole position. We advanced the electrode until there was visually responsive neuronal activity on most channels and until the local field potentials showed a distinct spectral signature of the cortical sheet, characterized by large amplitude alpha/beta (10-30 Hz) oscillations in deep channels and gamma (40-150 Hz) oscillations in superficial channels [132].

We used MonkeyLogic (developed and maintained at the National Institute for Mental Health [137]) to control the behavioral task. Visual stimuli were displayed using PROPixx Pro projectors (VPixx Technologies, Quebec, Canada) with a resolution of 1920 x 1080 at a 120 Hz refresh rate. The projector screen was positioned 57 cm in front of the monkey’s eyes. Luminance calibration was performed with a Photo Research PR-650 at the projector screen with a median value of 2.613 cd/m2. The monkey was trained to fixate its eyes around a central fixation dot (radius of fixation window: 1.5 visual degrees). Eye position was monitored using an Eyelink 1000 (SR Research, Ottawa, Canada). A task began with an isoluminant gray screen. Once the monkey maintained fixation for 500ms, a sequence of drifting gratings appeared (radius = 12 visual degrees, drift rate: 2Hz, 1 cycle/visual degree, 0.8 Michelson contrast, angle of drifting grating was 45 or 135 degrees from the vertical midline) for 500ms, and was replaced by a blank screen with the fixation dot remaining for 500ms. This sequence was repeated 4 times in a trial, for a total trial length of 4500ms (500ms pre-sequence fixation + 4×1000ms stimulus presentations). The monkey was rewarded for maintaining fixation throughout the trial with a small juice reward. Only the first 1000ms of data was used for this analysis.

## Analysis

Single units were sorted using Kilosort4. We only selected neurons which satisfied both quality requirements (Signal-to-noise ratio > 1.2, Presence ratio > 97.5%). We convolved all single unit spike times with a post-synaptic potential kernel with a rise time constant of 0.5ms and a decay time constant of 4.5ms. This convolution was applied to the model’s response and to the neurophysiological spike time data. This step was necessary in order to facilitate spectral analysis, which is more well-behaved on continuous data as opposed to point-process data [134].

We estimated each neuron’s spectrogram (time-frequency response) using multitaper spectral estimation with a Kaiser window of length 400 ms and 98% overlap between adjacent time windows. For both the model and the electrophysiology data, we performed this analysis from 500 ms before stimulus onset to 1000 ms after stimulus onset. Time-frequency responses during stimulus presentation (Fig. 6; Supplemental Figs. S16-S18 for Izhikevich analyses) were used as GSDR objectives for generating neural responses with similar spectral structure.

## Acknowledgements

We thank Nancy Kopell (and the Cognitive Rhythms and Cognition - CRC team), Homero Esmeraldo, Eli Sennesh for their critical comments to an initial version of this manuscript.

## Code availability

Github repository: https://github.com/DynaSim/DynaSim/tree/devDL https://github.com/HNXJ/GSDR

