## Supplemental file for "A genetic algorithm for self-supervised models of oscillatory neurodynamics"

**(Supporting information)**

**Code availability**: Github repository(s):

https://github.com/DynaSim/DynaSim/tree/devDL (Dynasim on MATLAB)

https://github.com/HNXJ/GSDR (Jax/Jaxley on Python)

### Supporting Information

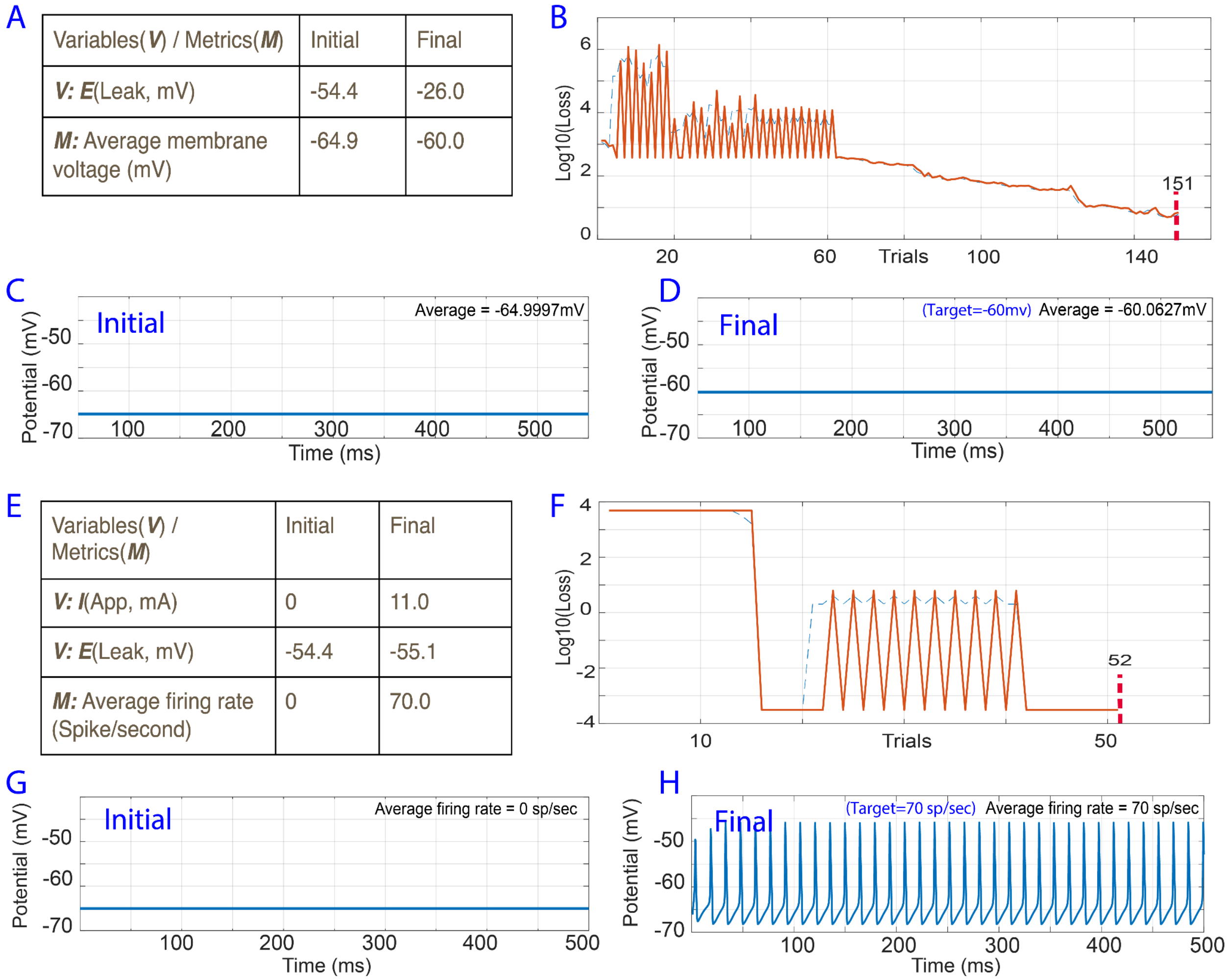

A

| Variables(*V*) / Metrics(*M*) | Initial | Final |
|---|---|---|
| ***V:*** ***E***(Leak, mV) | -54.4 | -26.0 |
| ***M:*** Average membrane voltage (mV) | -64.9 | -60.0 |

E

| Variables(*V*) / Metrics(*M*) | Initial | Final |
|---|---|---|
| ***V:*** ***I***(App, mA) | 0 | 11.0 |
| ***V:*** ***E***(Leak, mV) | -54.4 | -55.1 |
| ***M:*** Average firing rate (Spike/second) | 0 | 70.0 |



***Supplemental Fig. S1****: Single cell basic optimizations. (A) Summary of initial and final training variables and metrics (B) Loss in trials corresponding to the target metrics. (C) Initial membrane potential response. (D) Final membrane potential response. (E) Summary of initial and final training variables and metrics for firing rate optimization. In this simulation both the reversal potential of the leak channel (Eleak) and the internal drive (Iapp) are variables that change in training. (F) Loss in trials corresponding to the target metrics. (G) Neuron response (Initial) (H) Neuron response (final)*

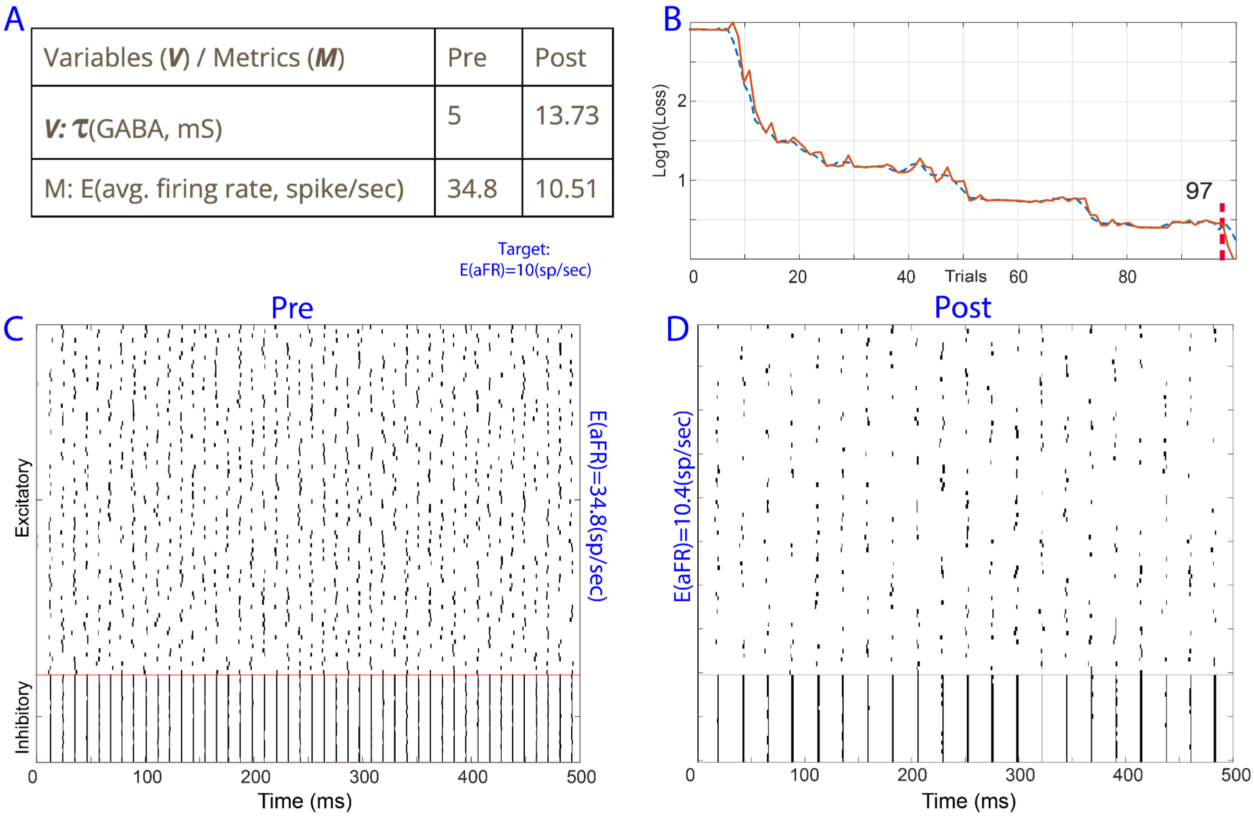


***Supplemental Fig. S2***: *E-I (Excitatory-Inhibitory) neuronal optimization of population firing rate and membrane potential. (A) Summary of initial and final training variables and metrics. (B) Loss in trials corresponding to the target metrics, firing rate and membrane potential. (C) Initial spiking response as a raster plot. Each row corresponds to a single neuron's spiking during one trial (500ms). (D) Raster plot after training.*

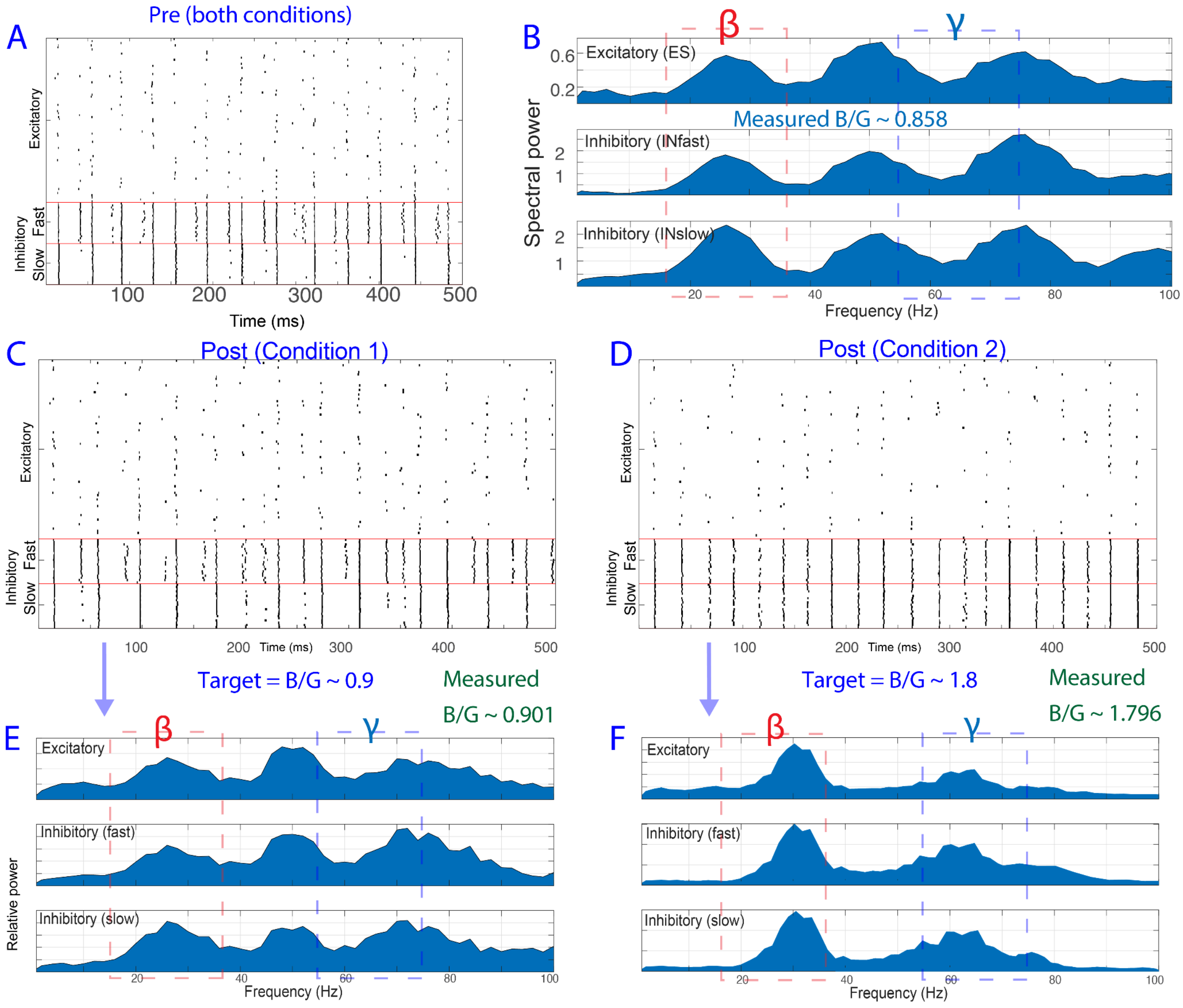


***Supplemental Fig. S3****: E-I dual-context specific band frequency ratio optimization responses before (A-B) and after training (C) Post-training raster plot, condition 1 (D) Post-training raster plot, condition 2 (E) Spectral response, condition 1. (F) Spectral response, condition 2.*

A

| Variables(***V***) / Metrics(***M***) | Pre | Post |
|---|---|---|
| ***V:*** τ(GABA, ms) | 5 | 8.05 |
| ***M:*** Avg. firing rate *(Exc., spike/sec)* | 34.83 | 19.61 |
| ***M:*** Avg. firing rate *(Inh., spike/sec)* | 91.81 | 71.41 |

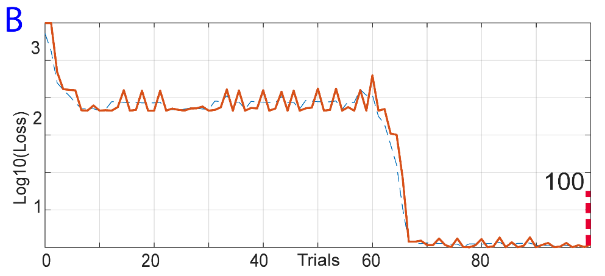


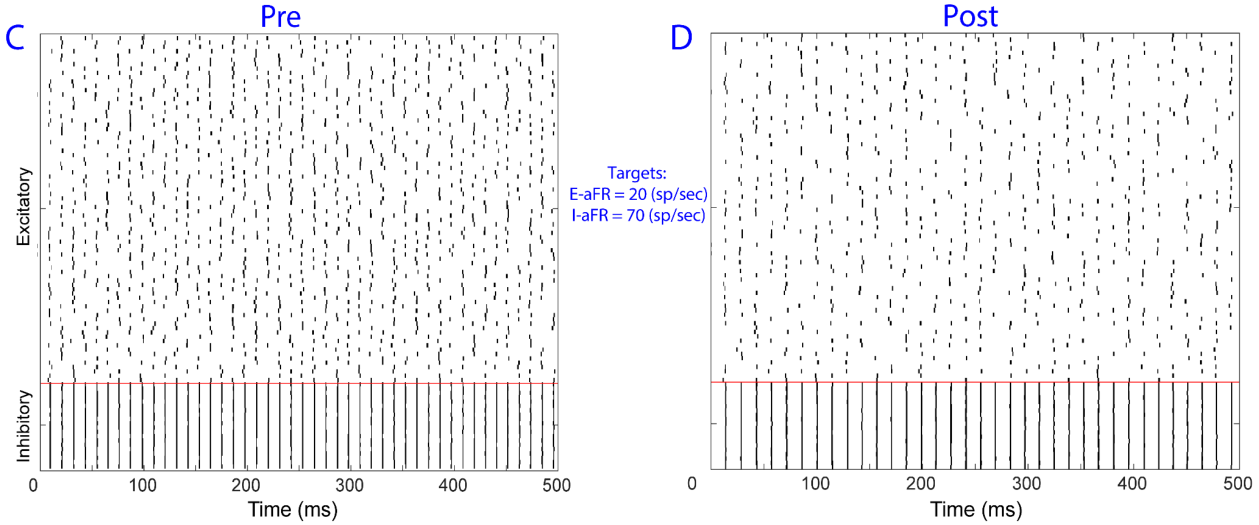


***Supplemental Fig. S4****: E-I (Excitatory-Inhibitory) neuronal population multitarget firing rate optimization. (A) Summary of initial and final training variables and metrics. In this simulation, the firing rates of E and I populations should converge to their respective objectives (Target firing rate of 70 spikes per second for I population and 20 for E population. (B) Loss in trials corresponding to the target metrics. Each population's firing rate converges to the desired target. (C) Initial raster plot. (D) Raster plot after about 100 trials.*

A

| Variables(*V*) / Metrics(*M*) | Pre | Post |
|---|---|---|
| *V:* τ(GABA, ms) | 5 | 29.98 |
| *V: Noise (Exc.)* | 40 | 50.00 |
| *V: Noise(Inh.)* | 40 | 41.86 |
| *M: Nat. freq. (Hz)* | 91.27 | 30.60 |

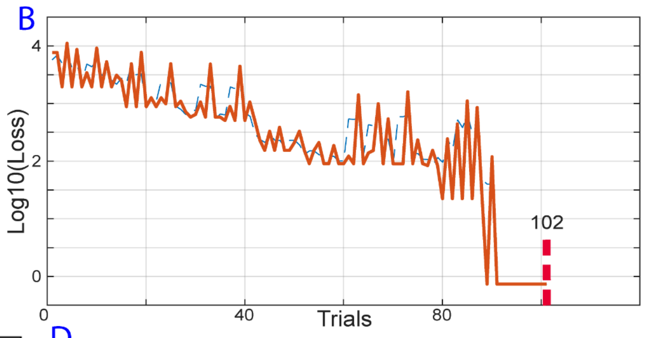


C

| Parameter log | Trial 1 | Trial 25 | Trial 50 | Trial 75 | Trial 102 |
|---|---|---|---|---|---|
| *V:* τ(GABA, ms) | 5 | 17.48 | 22.00 | 24.12 | 29.98 |
| *V: Noise (Exc.)* | 40 | 39.58 | 47.03 | 50.00 | 50.00 |
| *V: Noise(Inh.)* | 40 | 43.32 | 50.00 | 44.56 | 41.86 |
| *M: Nat. freq. (Hz)* | 91.27 | 75.65 | 64.27 | 54.85 | 30.60 |

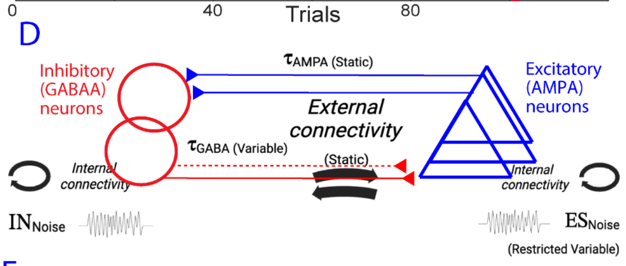


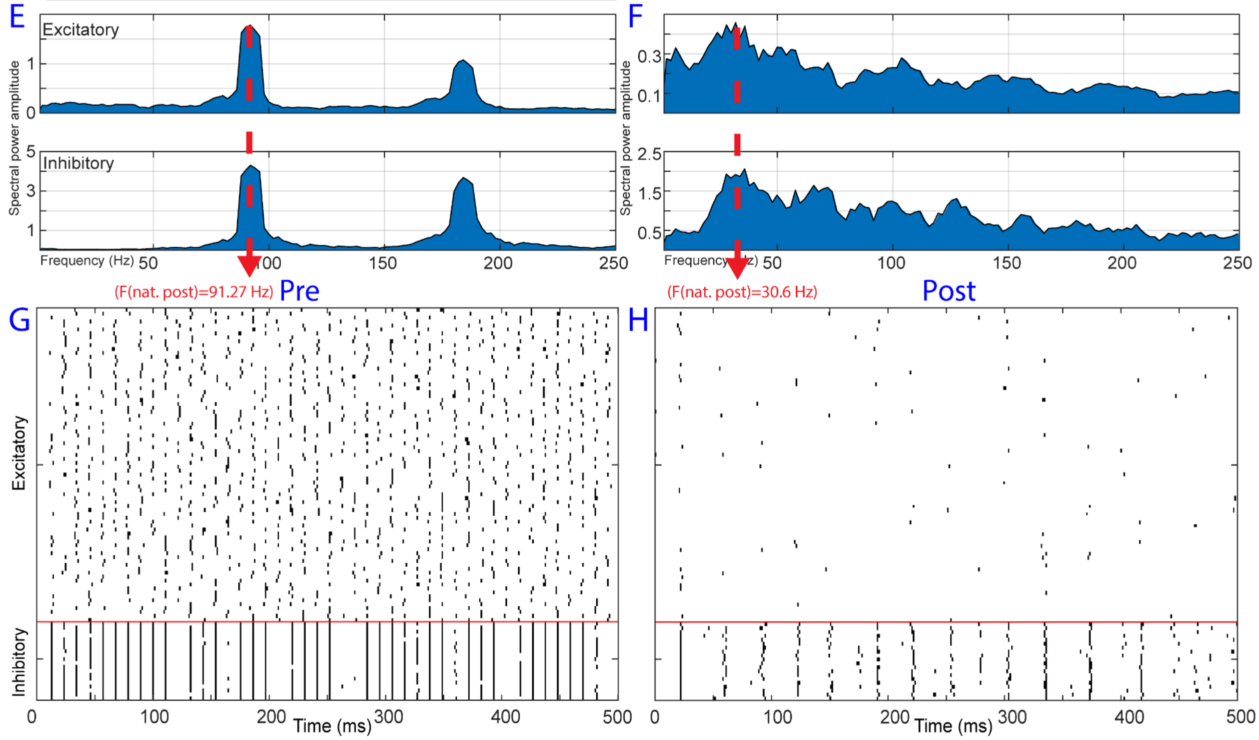


***Supplemental Fig. S5****: E-I natural (dominant) frequency tuning. (A) Summary of initial and final training variables and metrics. In this simulation time constant of inhibitory synaptic transmission and internal noise are optimized to achieve a natural (dominant) spectral peak at a particular frequency (30Hz). (B) Loss in trials corresponding to the target metrics. (C) Summary of training variables and metrics during training. (D) neural circuit schematic. (E) Initial frequency response. (F) Frequency response after training showing that the neural circuit has achieved the target metrics. (G) Initial raster plot. (H) Final raster plot.*

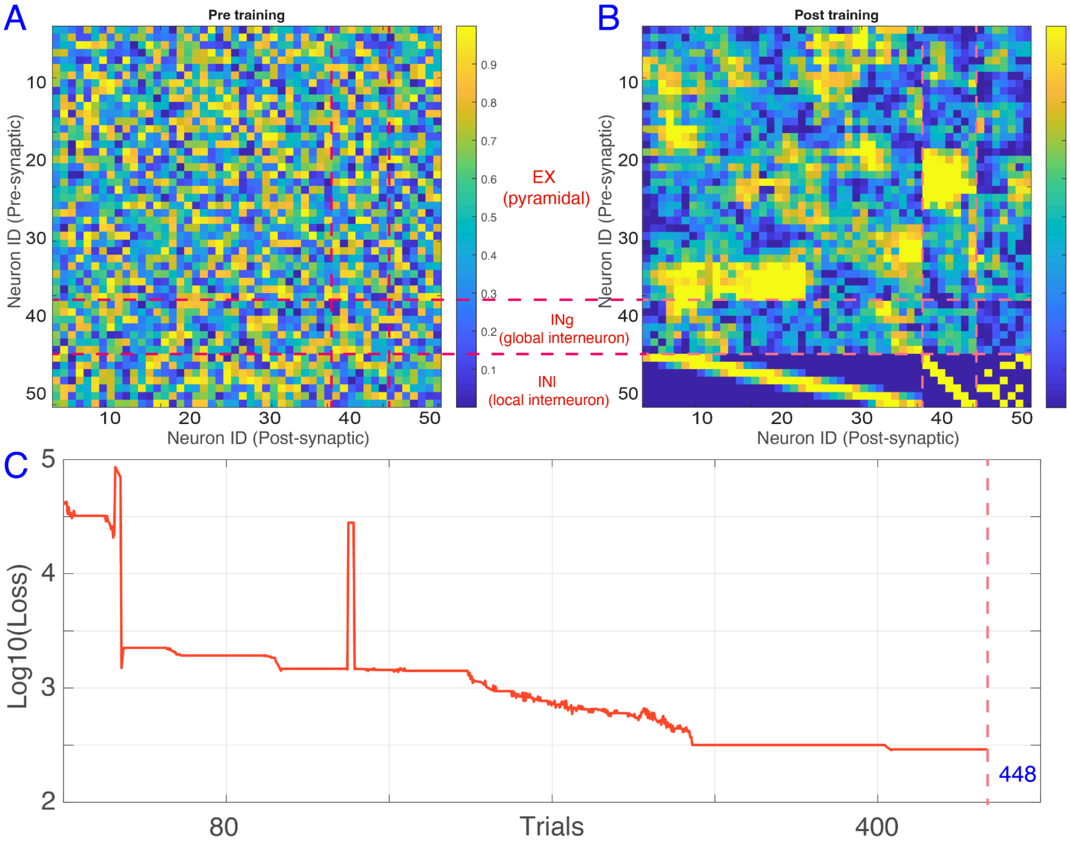


***Supplemental Fig. S6:*** *(A-B) Pre-Post training synaptic connectivity matrices, corresponding to Fig. 6. The model formed asymmetric connections between neurons, such that it resulted in the stimulus-induced gamma. Notably, local-interneurons formed sparse and strong connections, instead of distributed and weak synapses. In addition, several interconnected sub-networks (E-E, E-INg, E-INl) have been formed. (C) Log-space (base of 10) loss across trials during training for the model in Fig. 6, up to trial 448 (last trial)*

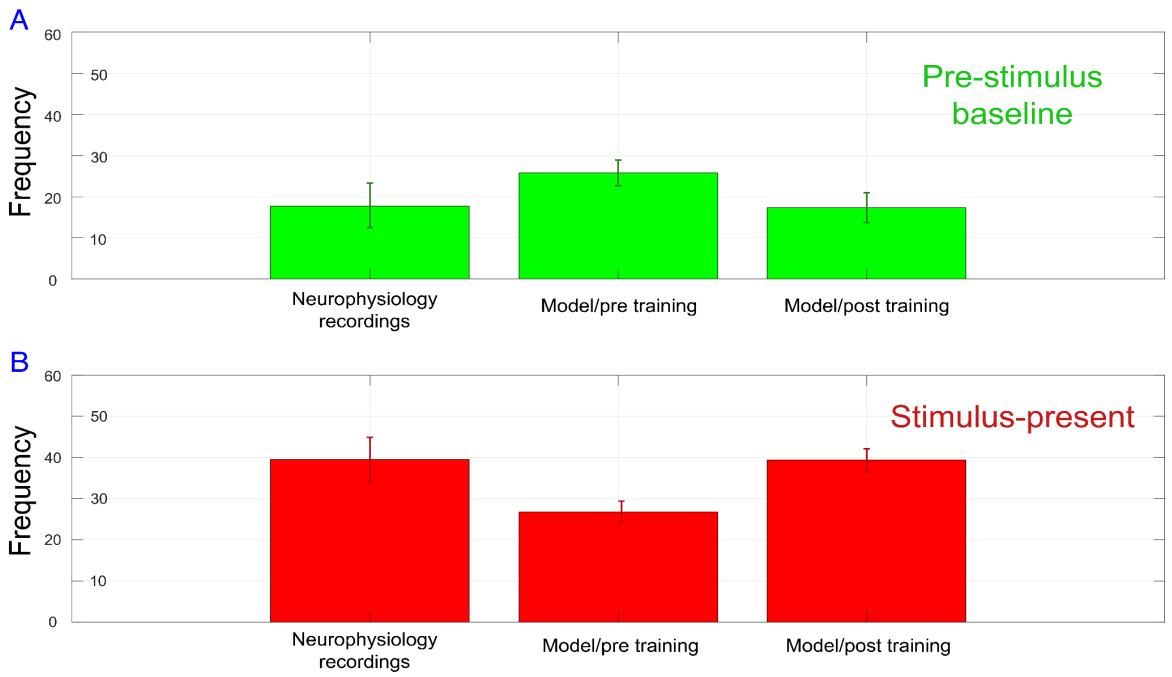


***Supplemental Fig. S7**: Dominant frequency in the recorded spiking and model (pre training/post training, pre-stim/post-stim). The upper subpanel corresponds to the pre-stimulus baseline and bottom subpanel corresponds to the stimulus-present time. Error bars are across number of neurons (N = 80 neurons in recording, N=50 neurons in the model)*

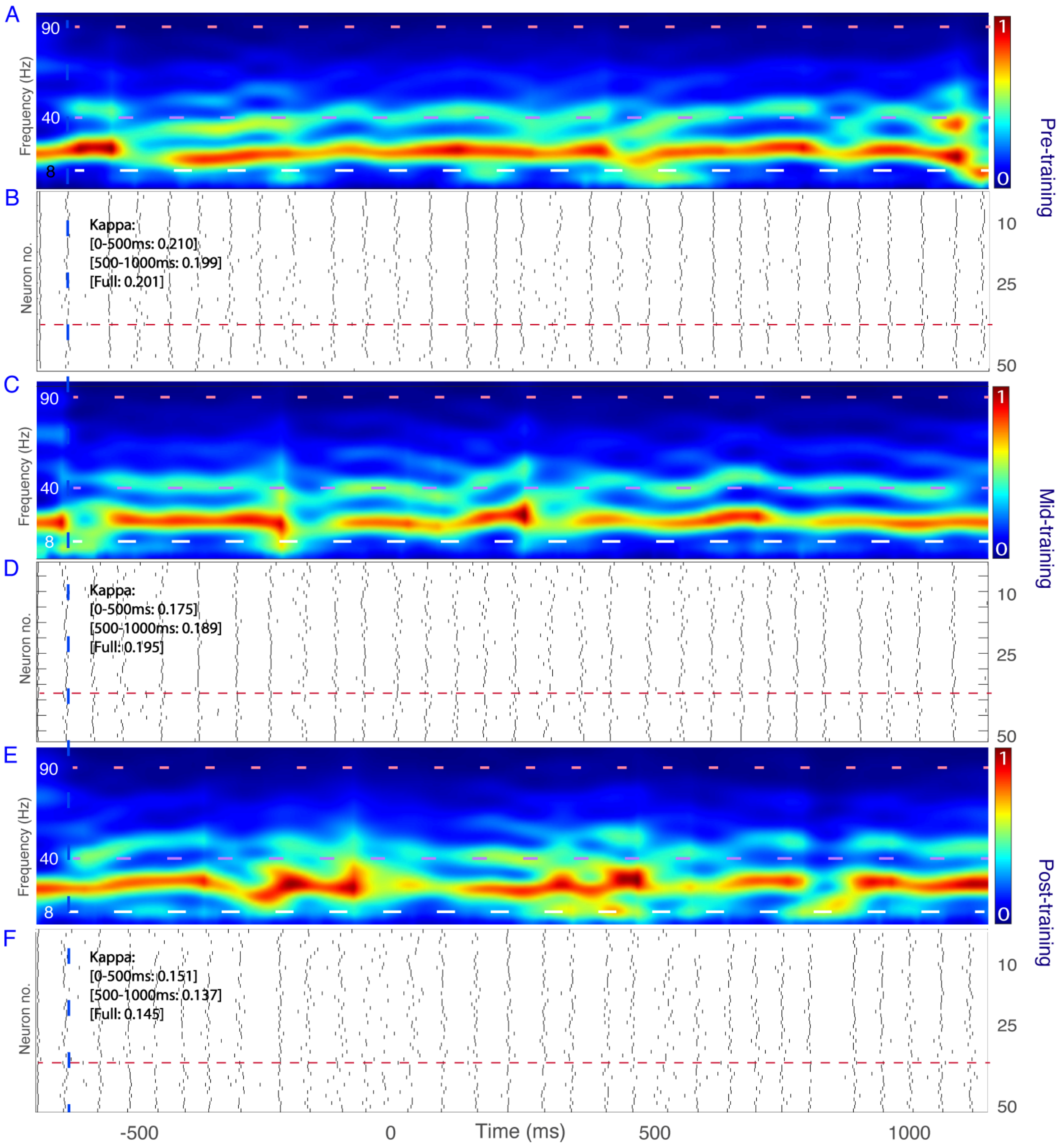


***Supplemental Fig. S8****: Single unit neural baseline responses from the model. (A) Time-frequency response (scaled to 0 as the lowest power and 1 for the highest power, 1/f adjusted) of the model , pre-training (B) Corresponding raster plot. (C) Time-frequency response of the model , mid-training (D) Corresponding raster plot, mid-training. (E) Time-frequency response of the model , post-training (F) Corresponding raster plot, post-training.*

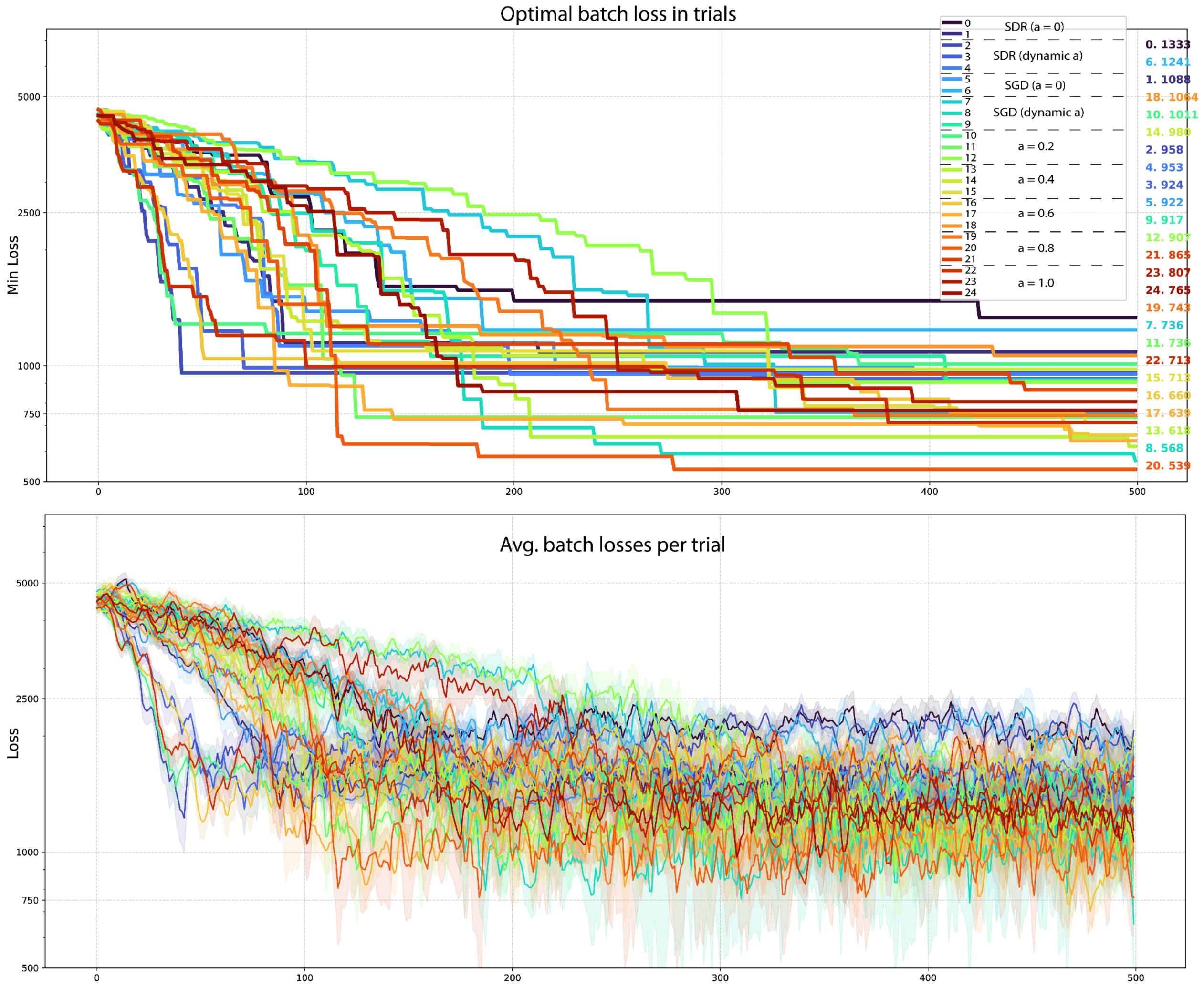


***Supplemental Fig. S9****: Comparison of different GSDR self-supervision models on the spectral similarity task. All models used here had identical initial parameters and ran for 500 trials (plotted on the x-axis) for evaluation. The value of the loss function (log-ratio spectral loss between the PSD of model and data, see Equation 4) on the spectral similarity task is plotted on the y-axis. (upper subplot) Optimal loss up to each trial. (lower subplot) The average and standard deviation of the batch (size = 4) at each trial. When alpha=0 (models with index 0, 1, 5, 6, see Legend on upper subpanel), there is no self-supervision and GSDR only follows delta rule (SDR or SGD depending on the setting). At alpha=1.0 (models with index 22, 23, 24), supervision has no effect (delta gets ignored). When alpha is dynamic or non-zero and non-one (models with index 2, 3, 4, 7-21), the models are self-supervised and are less likely to get stuck in a local minimum.*

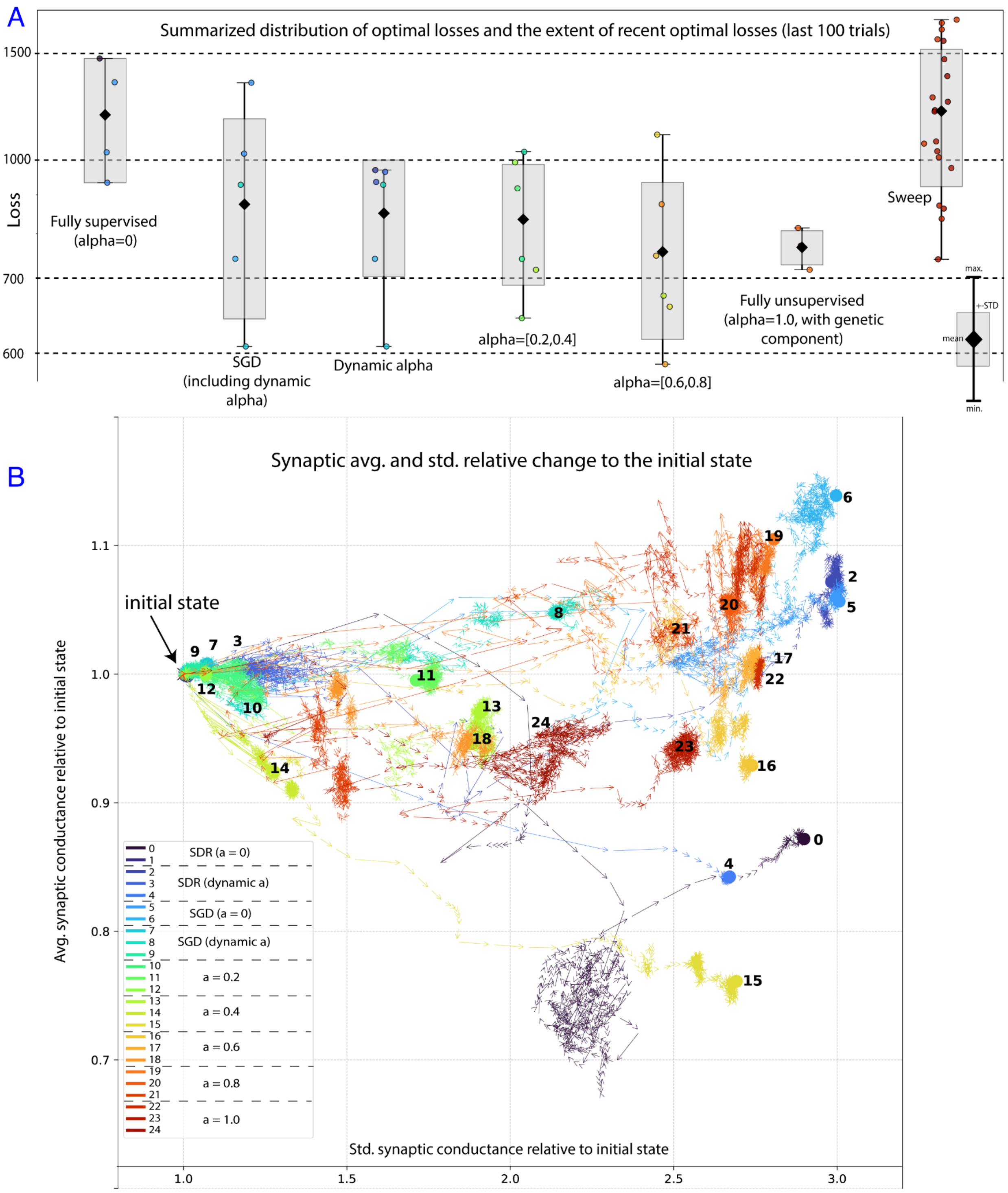


***Supplemental Fig. S10****: Comparison of different GSDR self-supervision models on the spectral similarity task, showing the average of the last 100 trials for each model type (annotated for each group). (A) Mean, standard deviation, extent (maximum and minimum), and the optimal point of each model are plotted per group, including across parameter sweep iterations (annotated as sweep). This comparison shows that self-supervision (represented by non-zero alpha) improves convergence to the objective since the similarity score between model and data is the worst when self-supervision is off (when alpha = 0). (B) Relative parameter average and standard deviation changes relative to initial state (1.0,1.0) across trials. This shows the change in parameters for each*

*model type, showing that all models had a comparable exploration (similar order of magnitude) of the parameter space.*

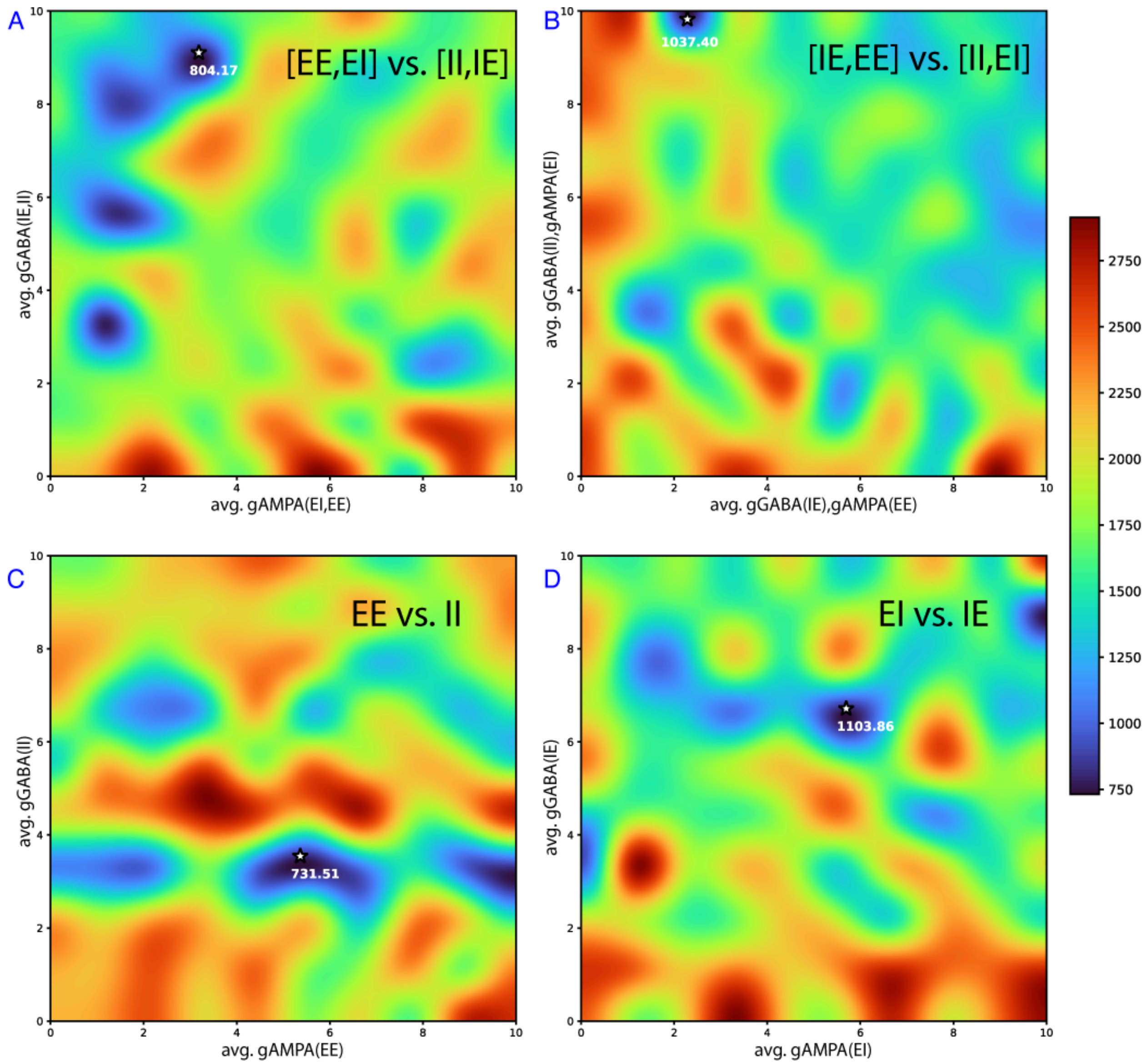


***Supplemental Fig. S11****: Parameter sweep across four groups of two-parameter set combinations. In each group, synaptic conductance weights are set to a uniform distribution centered at values 0-10 (x and y axis) and +- 0.5. For unassigned parameters in sweep, uniform random values in range (4.5,5.5) are used. Parameter group pairs are swept across values 0-10 (10 rows, 10 columns) with 10 repetitions (4,000 total runs per subpanel, equivalent to a 500 trial optimization with batch size of 4, approximately twice the total computational cost of*

*each model in Supplemental Fig. S9 & 10). The heatmap shows the minimum instance of loss across the 10 repetitions for each two-parameter combination. (A) all AMPA vs. GABA synapses. (B) All synapses in which the post-synaptic neuron is E vs I. (C) All EE vs. II synapses. (D) All synapses in which pre and post synaptic neurons are not from the same group.*

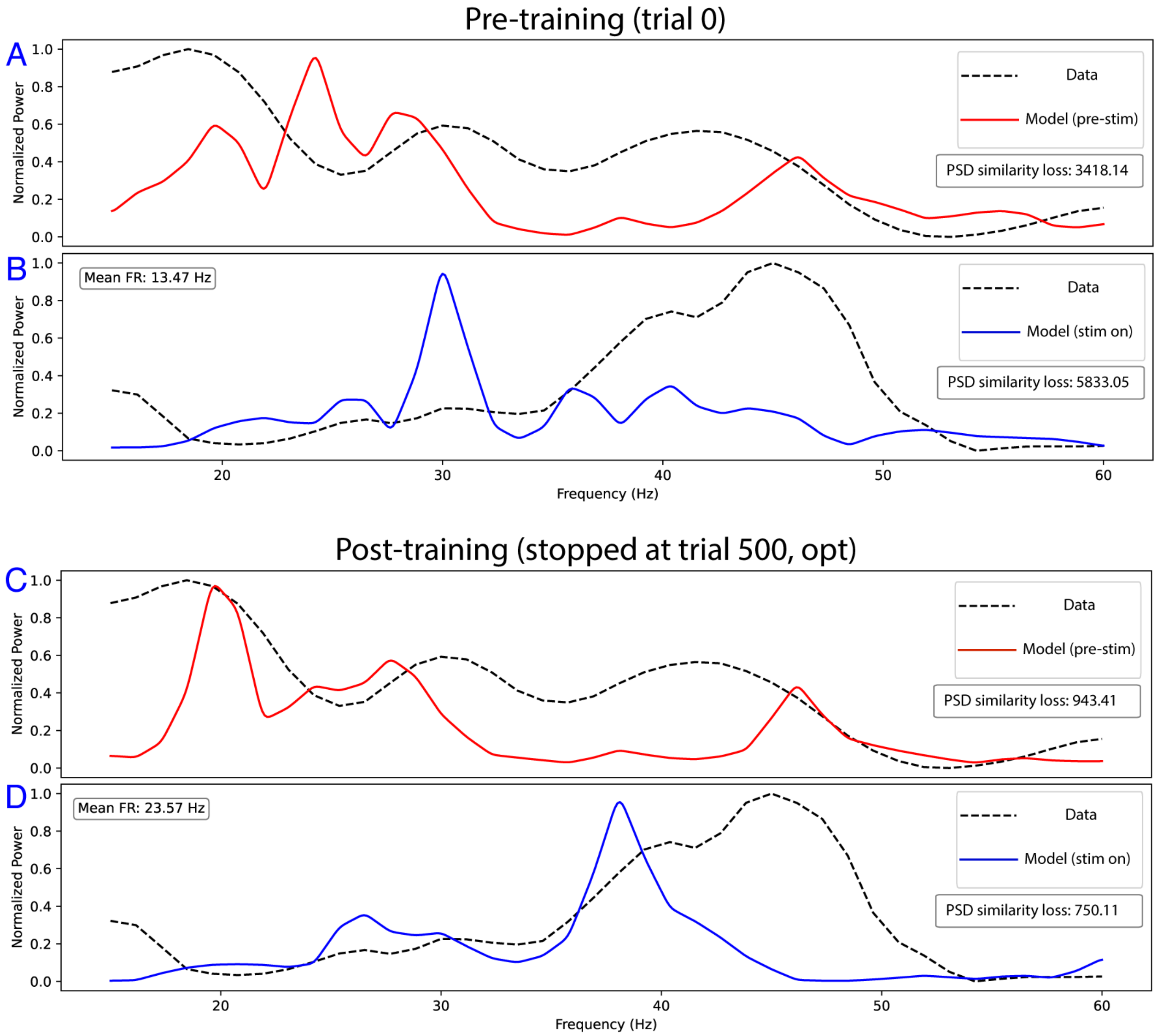


***Supplemental Fig. S12****: Power Spectral Density (PSD) example for models (solid lines) vs. data (dashed lines). The PSD (plotted on y-axis) was scaled by the maximum power across time and frequency (x-axis), separately for the pre-stimulus and stimulus periods. The normalized PSD is plotted at trial 0 (initial state, A-B) and after 500 trials (C-D) trained with GSDR (with dynamic self-supervision) on the spectral similarity task. The model-data spectral similarity loss is noted for each subpanel. As the training progresses, the model's PSD is pushed towards the PSD of the data for both the pre-stimulus period (shown in red, which tends towards the alpha-beta frequency band) and the stimulus period (shown in blue, which tends towards the faster gamma frequency band). Note that the pre-stimulus PSD of the data has a peak in the beta range ~18Hz which is not apparent in the Time-Frequency Representation of the same data (Fig. 5C) because the normalization in Fig. 5C was performed on the entire duration of the signal.*

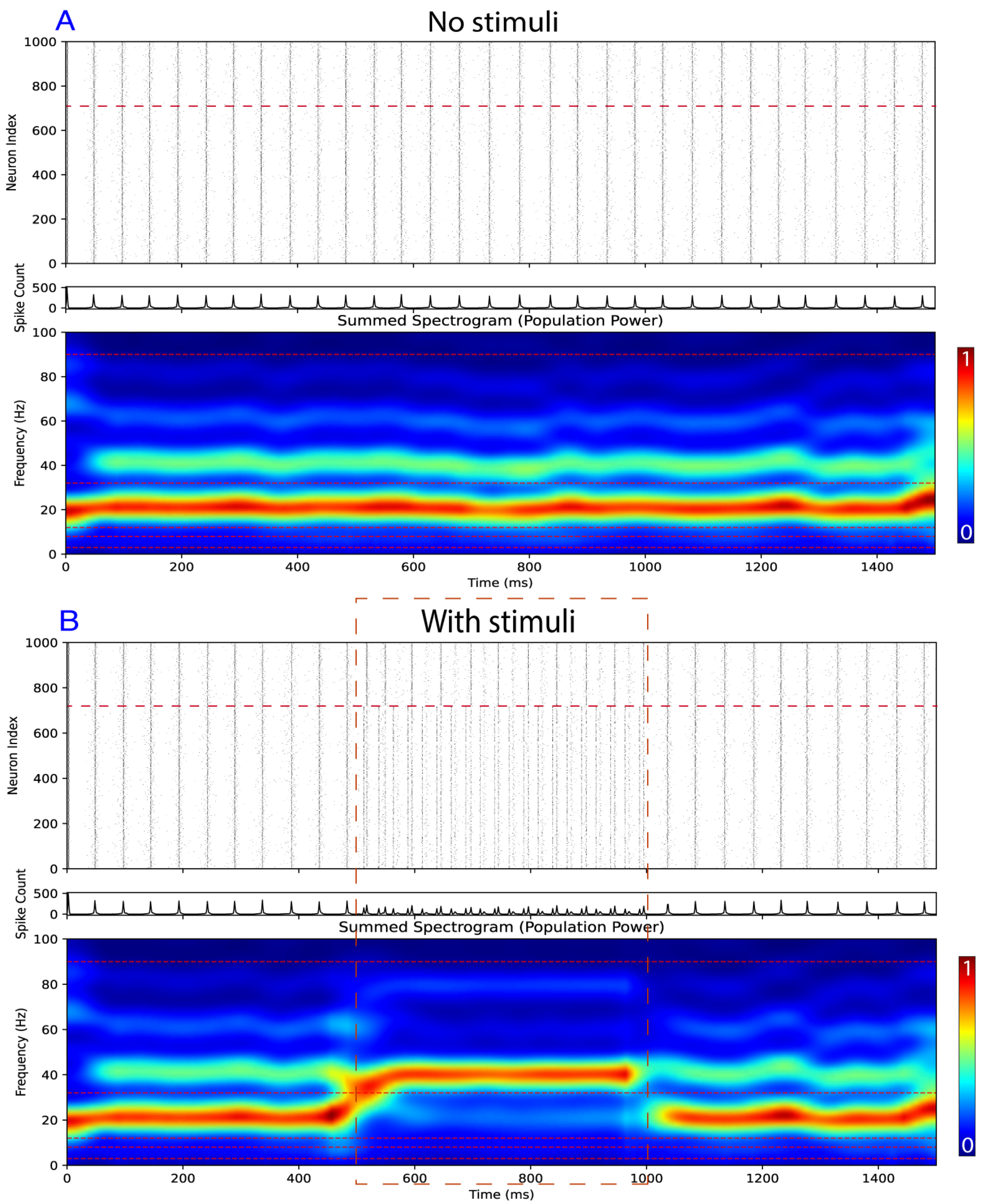


***Supplemental Fig. S13****: Time Frequency Response (normalized) for augmented model. Augmentation process is performed via upsampling and gaussian smoothing with kernel size equal to the upsampling scale. For this model, the upsampling scale was 20 (50 neurons projected to 1000 neurons). All neuronal parameter arrays are either 1-dimensional (i.e., one per neuron for Hodgkin-Huxley parameters) or 2-dimensional (i.e., one per pair of pre-post synaptic neurons for gAMPA, gGABA… ). each array is upsampled proportional to the upsampling scale and smoothed (similar to upsampling/zoom on images). The augmented model contains E cells in index 1-720 (1-36 in original model) and I cells in index 721-1000 (37-50 in original model). The augmented model engages in a weak-PING, similar to the original model.*

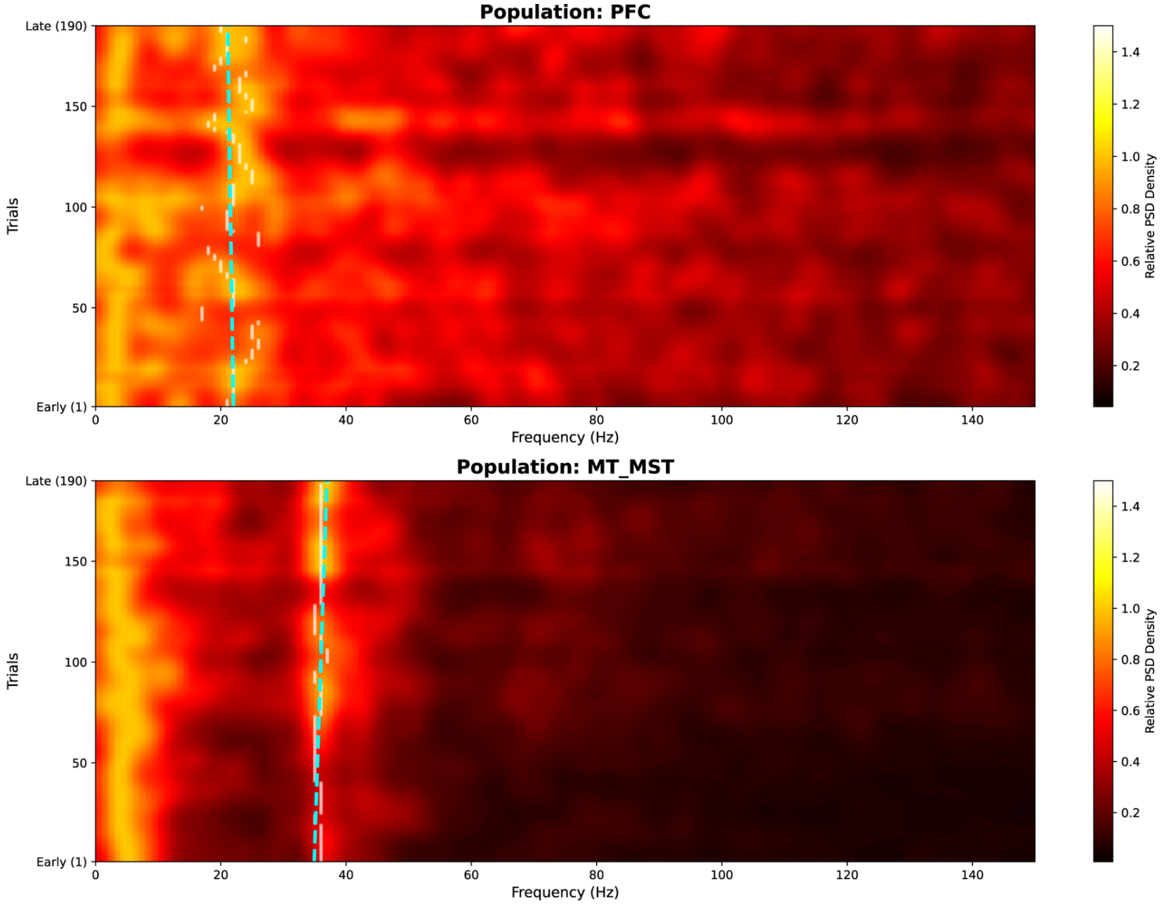


| Area | N Units | $R^2$ | p-val | Slope | Shift | MSE | K-Base | K-Stim |
|---|---|---|---|---|---|---|---|---|
| PFC | 122 | 0.019 | 5.85e-02 | -0.0045 | -0.86Hz | 3.2155 | -0.0003 | -0.0004 |
| MT_MST | 95 | 0.136 | 1.65e-07 | 0.0132 | +2.50Hz | 3.3522 | 0.0062 | 0.0055 |

***Supplemental Fig. S14****: Trial-by-trial power spectrum density (PSD) of single neuron signals. Both subpanels are from the same session, simultaneous recordings during identical trials. The analysis was performed 500ms before the onset of a visual stimulus until 500ms after the offset (total duration = 1500ms, n=190 trials). The upper subpanel shows the PSD of the prefrontal cortex (n=122 single neurons). The lower subpanel shows the PSD of the area MT/MST (n=95 single neurons). The gamma frequency (~38-40Hz) of area MT/MST is observed to increase by ~2.5Hz across trials, as previously reported (Brunet et al., 2014 [1]). However, this gamma oscillation is absent in the simultaneously recorded PFC signal (upper subpanel). The PFC signal instead has a peak frequency in the beta band (~20Hz), consistent with previous reports in the LFP signal (Lundqvist et al., 2020 [2]). Unlike potential artifacts or noise sources which would be either consistent across trials (i.e., the slope of electric notch noise such as the line or screen noise at 60Hz would be absolute 0) or identical across nearby probes (external electromagnetic noise appears with the same spectral profile across all probes). However, both the slope and peak frequency are different across areas, showing natural, non-static neuronal dynamics.*

synchrony (Kappa) in strong pyramidal-interneuron gamma network (PING) models with different baseline noise amplitude

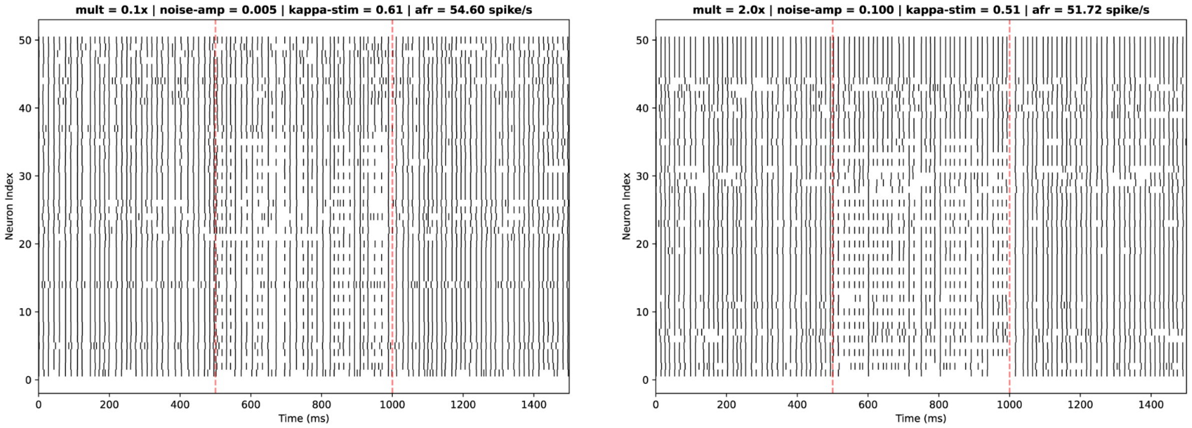


***Supplemental Fig. S15****: Synchrony in strong-PING model. Notice that the kappa value (Rook and Penning, 1991) for hypersynchronized activity is ~5 to ~20 times higher than the simulation in Fig. 6 (and Supplemental Fig. S6). Despite the presence of internal noise, the synchrony is still order of magnitude (~ 0.5 compared to ~0.05) stronger than simulations (Fig. 6, Supplemental Fig. S8) and data (Fig. 5, Supplemental Fig. S14)*

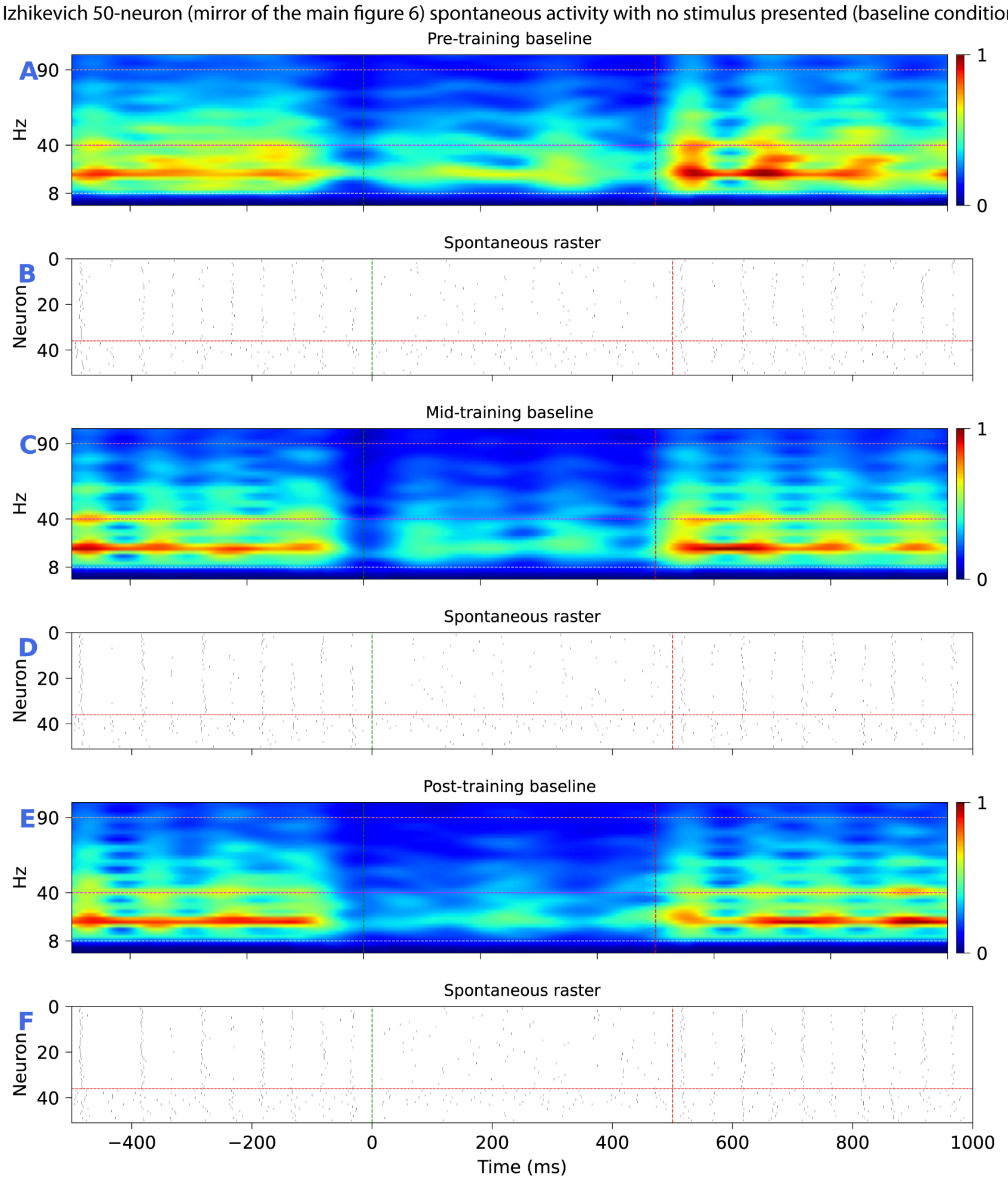


***Supplemental Fig. S16***: *Izhikevich reduced-spiking baseline simulation.*

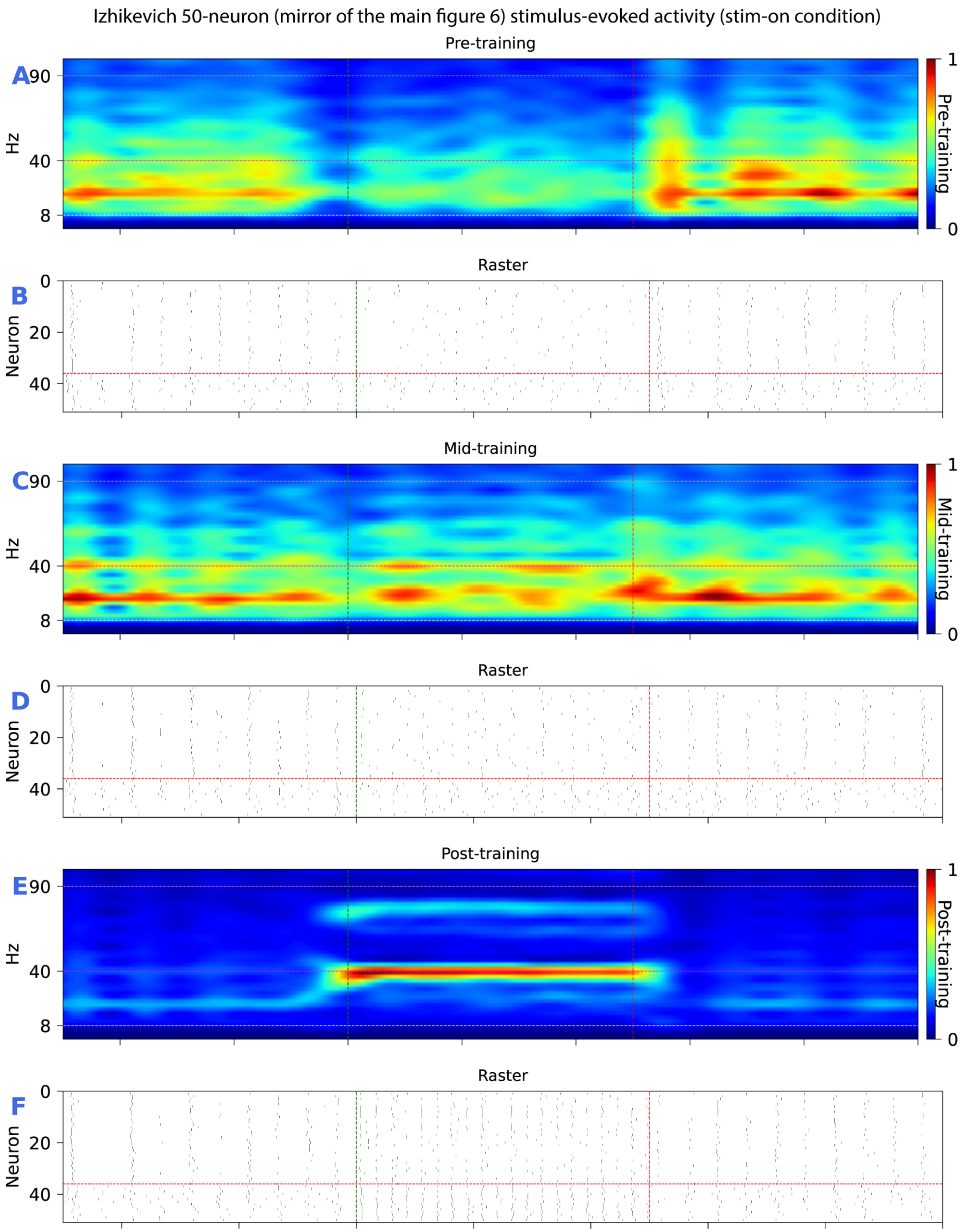


***Supplemental Fig. S17****: Izhikevich reduced-spiking stimulus-on simulation.*

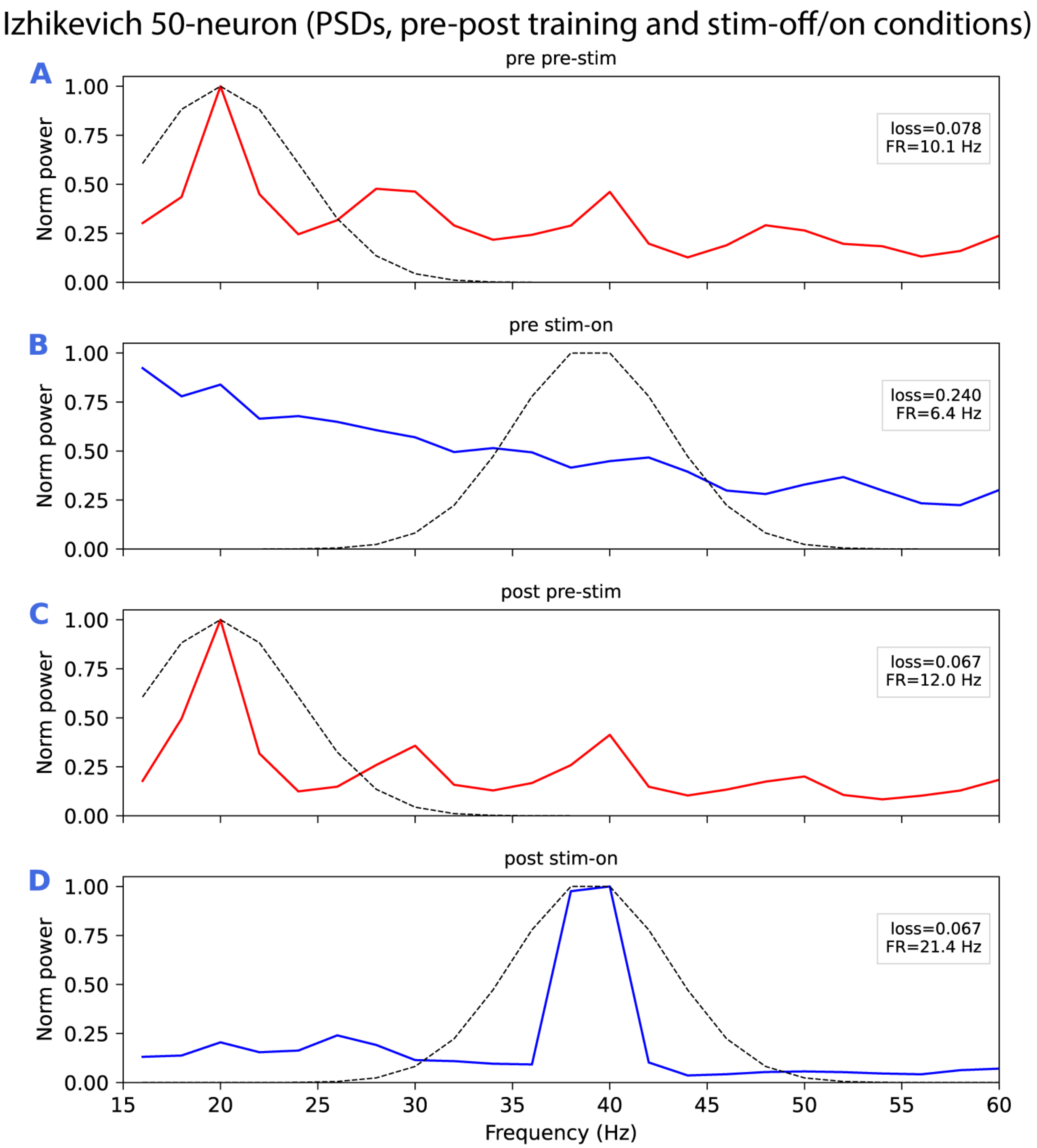


***Supplemental Fig. S18**: Izhikevich reduced-spiking PSD comparison across baseline and stimulus-on conditions.*

### *Supplementary text*

**Synaptic time constants in excitatory-inhibitory (E-I) networks:** After establishing good performance with this toy optimization problem and other more basic scenarios in neuronal circuits (see Methods and Supplemental Fig. 1 for simple single neuron optimization problems and Supplemental Fig. 2 for optimizing population firing rates), we next tested our approach in a multi-objective optimization problem by setting two firing rate objectives for the population of excitatory (E) and inhibitory (I) cells. We implemented an excitatory-inhibitory (E-I) model comprising 100 neurons, including 80 excitatory and 20 inhibitory neurons (Supplemental Fig. 4). This model incorporated specific mechanisms: AMPA receptors for excitatory synapses and GABA receptors for inhibitory synapses, along with other parameters influencing connectivity such as synaptic connections, synaptic time constants, and both excitatory (E-noise) and inhibitory (I-noise) noise. Initial synaptic connectivity was established using a uniform matrix across all neurons. We trained the model to pursue two parallel objectives with equal weight, aiming for average firing rates of 20 spikes/sec for excitatory neurons and 70 spikes/sec for inhibitory neurons. Initially, the excitatory and inhibitory populations exhibited firing rates of 34.8 spikes/sec and 91.81 spikes/sec, respectively (Supplemental Fig. 3A). Only the inhibitory synaptic time constants were permitted to change. As illustrated in Fig. 3, the model successfully met the target firing rates by increasing the inhibitory time constants, showcasing the capacity of GSDR in modulating mechanistic parameters to simultaneously optimize multiple objectives.

Although synaptic time constants and firing rate are inversely correlated, their relation is not perfectly linear (i.e, multiplying the time constant by 2 will not necessarily divide the frequency by 2). In the model (Fig. 3), the ratio of time constant (post to pre) is ~1.6 but the corresponding ratio for average firing rates are different (1.77 for the excitatory population, 1.28 for the inhibitory population). This sort of nonlinear interactions makes it difficult to predict solutions and manually tune biophysical models to reproduce desired responses in multi-objective cases.

**Population natural frequency with noise and synaptic time constant**: Next, we used the same neuronal model implemented in Fig. 3 to tune the natural frequency of its average neuronal potential. The natural frequency is defined as the frequency with the maximum oscillatory power in a neural circuit (Fig. 4E-F). The pre-training natural frequency of the excitatory population was 91Hz (Fig. 4A). For training, inhibitory synaptic time constants and baseline Gaussian noise gain were allowed to vary in the model. As seen in Fig. 4, the model converged to the target natural frequency (30Hz) by increasing inhibitory time constants. The results indicate that the relation between synaptic time constants and spectral response of the model is negatively correlated yet pre to post time constant ratio and the natural frequency peak are not linearly related (Fig. 4A,C). We also note that the model achieved the objective by creating a highly inhibited state for the excitatory E cells (Fig. 4H), consistent with an ING mechanism, where a neuronal oscillation is sustained by the I cells. Note that no other objective other than the peak frequency of the model was specified, but in principle, other objectives can be explicitly stated (e.g., to maintain E cell firing above a particular threshold) if desired. It is interesting that the model has converged to a beta rhythm that leads to overall more network inhibition (amongst the E cells), a result consistent with the suggestion of Predictive Routing that beta states are associated with prediction-driven inhibition.

The tradeoff between the unsupervised and supervised terms is controlled by alpha, ranging from 0 to 1. Alpha = 0 corresponds to a fully supervised update, whereas alpha = 1 corresponds to an activity-dependent update with stochastic exploration. In dynamic-alpha models, alpha follows a bounded stochastic random walk and is filtered by the same genetic selection/deselection logic as other parameters. Comparative benchmarking on the spectral-similarity task demonstrated that hybrid self-supervised GSDR, especially intermediate alpha values and dynamic alpha, produced better average performance than fully supervised models (Supplemental Figs. 9 and 10) and parameter-sweep baselines (Supplemental Fig. 11) under matched constraints.
